# Wildfire alters nitrogen cycling to increase soil emissions of nitric oxide (NO) and the heterogeneity of nitrous oxide (N_2_O) in California chaparral

**DOI:** 10.1101/2025.09.22.675225

**Authors:** Elizah Z. Stephens, M. Fabiola Pulido Barriga, Aral C. Greene, Alexander H. Krichels, Meg Kargul, Loralee Larios, Sydney I. Glassman, Peter M. Homyak

**Author notes:** **Corresponding author: Elizah Z. Stephens**, Department of Environmental Sciences, University of California, Riverside CA 92521, USA.

## Abstract

Wildfires can disrupt ecosystem nitrogen (N) cycling by combusting vegetation biomass N and depositing ash that is rich in ammonium (NH_4_^+^) onto soils. Post-fire increases in NH_4_^+^ and soil physicochemical changes may promote further N loss by stimulating microbially-driven emissions of nitric oxide (NO) and nitrous oxide (N_2_O)—trace gases that alter air quality and climate. We hypothesized that soil NO and N_2_O emissions would increase with the flush of post-fire N availability and would be highest in soils that burned at medium severity due to the combination of high soil N availability and persistence of microbial activity. To test this, we established nine plots (6 burned; 3 unburned) and sampled soils seasonally over three years after wildfire in a chaparral shrubland in Southern California, USA. Wildfire significantly increased soil extractable NH_4_^+^ by an average of 10 µg NH_4_^+^-N g soil^-1^, soil extractable NO_3_^-^ by 8 µg NO_3_^-^-N, and soil pH by 0.5 units. Post-fire soil NO emissions significantly increased by an average 74 ng NO-N g^-1^ (cumulative 40-h incubations) over three years, with the highest emissions measured in year one from plots that burned at medium and high severities. No significant effects of burning were detectable for N_2_O emissions over three years (average ± standard error; 224 ± 107 ng N_2_O-N g^-1^ soil in burned plots and 30 ± 17 ng N_2_O-N g^-1^ soil in unburned plots); however, we observed high fluxes only from soils that burned at medium and high severities (N_2_O > 3500 ng N_2_O-N g^-1^ soil). Isotopic characterization of N_2_O from high-emitting soils indicated contributions from diverse sources and increased N_2_O reduction to N_2_. Overall, the occurrence of high N_2_O fluxes and accelerated NO emissions post fire indicate wildfires interact with soil N cycling to promote burn-severity-sensitive gaseous N losses long after wildfires are extinguished.

## 1. Introduction

As global climate change exacerbates droughts and lengthens fire seasons, wildfires are projected to become more frequent and severe across many ecoregions globally (Ellis et al., 2022; IPCC, 2023; Sayedi et al., 2024). Increased fire frequency and severity can disrupt biogeochemical cycles and challenge the capacity of ecosystems to retain essential nutrients like nitrogen (N) (Dannenmann et al., 2018; Hanan et al., 2017; Underwood et al., 2018). The effects of wildfire on nutrient cycling may be particularly severe in ecosystems that are traditionally considered N-limited and exist in disproportionately fire-prone regions, such as dry shrublands (Stephens & Homyak, 2023; Underwood et al., 2018; Williams et al., 2019). Shrubland fires can volatilize approximately 60 % of vegetation-bound N directly to the atmosphere during combustion, transforming the remainder into ash rich in ammonium (NH_4_^+^; Dannenmann et al., 2018). With fewer plants to retain post-fire N as biomass, nutrient-rich ash is vulnerable to further losses from wind, run-off, and leaching (Goodridge et al., 2018; Guilinger et al., 2020; Hanan et al., 2016). Additionally, modeled hydrologic N losses in recently burned shrublands only account for about half of total N losses from the post-fire soil N pool (Goodridge et al., 2018), suggesting that significant N losses may also occur through gaseous pathways such as soil emissions of nitric oxide (NO), nitrous oxide (N_2_O), and dinitrogen gas (N_2_). NO and N_2_O are both highly relevant to global climate modeling as NO is a precursor for tropospheric O_3_ (Crutzen, 1979; Ostro et al., 2006) and N_2_O is a powerful greenhouse gas that destroys stratospheric ozone (Griffis et al., 2017; Ravishankara et al., 2009). Thus, post-fire soil NO and N_2_O emissions in shrublands may have significant but under-explored impacts on regional air quality and global climate feedbacks.

Dryland soils can produce large emission pulses of NO and N_2_O after wetting events when microbes rapidly metabolize N that has built up over the dry season (Homyak et al., 2016; Krichels et al., 2022, 2024; Leitner et al., 2017). Thus, fires that burn during dry periods may further prime soils for large pulses of gaseous N by providing excess N to surviving microbial communities (Krichels et al., 2022, 2024; Zhao et al., 2025). Previous studies in dry shrublands found that combined soil emissions of NO, N_2_O, and N_2_ over one year equaled ∼15% of the initial combustion losses of N, with relatively little N loss from leaching (Dannenmann et al., 2018). Similarly, studies in California chaparral identified up to 300% increases in NO emissions upon wetting burned soils and estimated that gaseous losses constituted 75% of total N loss from topsoils over 6 months post fire (Anderson et al., 1988; Anderson & Poth, 1989; Levine et al., 1988). Therefore, dry shrublands may be especially positioned to emit NO and N_2_O after fires, representing N loss pathways that are unaccounted for in the most comprehensive post-fire N budgets in chaparral to date (Goodridge et al., 2018; Hanan et al., 2016, 2017).

Soil NO and N_2_O emissions are largely produced as byproducts during microbial nitrification and denitrification (**Figure 1**). During nitrification, ammonia (NH_3_; measured as NH_4_^+^ in soils) is oxidized by nitrifying microbes to nitrate (NO_3_^-^) under oxic conditions (Firestone & Davidson, 1989), primarily by a phylogenetically constrained group of obligate chemoautotrophic Proteobacteria (ammonia-oxidizing bacteria; AOB) and Thaumarchaetoa (ammonia-oxidizing archaea; AOA; Hayatsu et al., 2008). During denitrification—a phylogenetically broad facultative process carried out by bacteria, archaea, and fungi—NO_3_^-^ is used as an alternative electron acceptor to oxygen in suboxic soil environments and can be sequentially reduced to N_2_ (Knowles, 1982). Despite significant reduction in overall microbial biomass by ∼45% after wildfire (Pressler et al., 2019), nitrification rates could be accelerated if significant NH_4_^+^ is deposited onto soils as ash (Karhu et al., 2015), and denitrification may be stimulated by the NO_3_^-^ generated by nitrification (Liu et al., 2015). Furthermore, high temperatures during combustion can generate cations and oxides which can elevate soil pH (Ulery et al., 2017) and form hydrophobic layers that lower water infiltration, keeping soils dry and aerobic (DeBano, 2000). Aerobic soil conditions may favor nitrification and higher soil pH and NH_4_^+^ may promote the activity of AOB over AOA, which are linked to higher NO and N_2_O production (Mushinski et al., 2019; Prosser et al., 2019). Elevated N_2_ production has also been measured in post-fire environments (Dannenmann et al., 2011, 2018), suggesting that high rates of denitrification may be promoted by soil physicochemical factors regardless of loss of microbial biomass.

**Figure 1.**
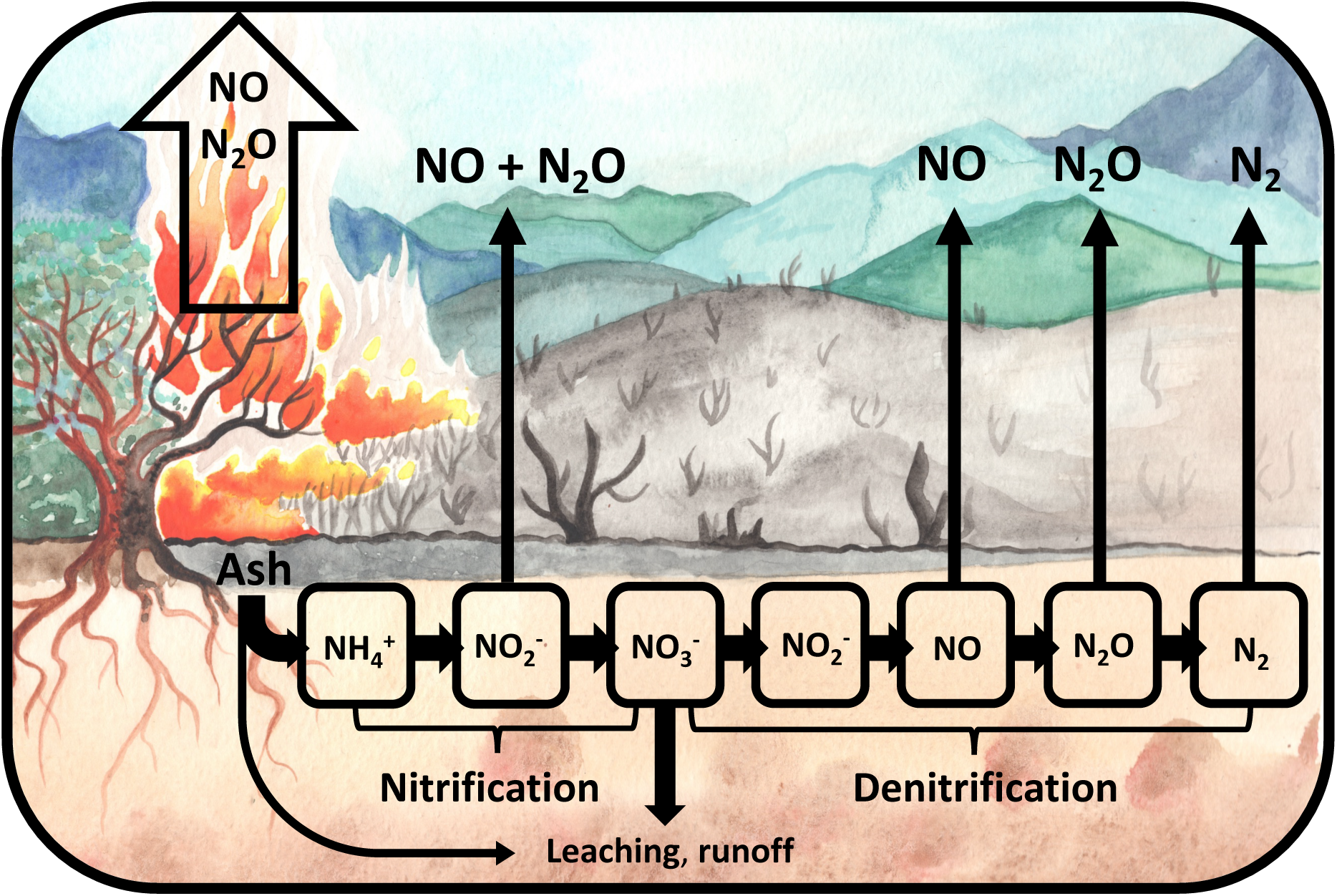
Sources of ecosystems N loss can include direct emissions of nitric oxide (NO) and nitrous oxide (N_2_O) to the atmosphere during combustion, post-fire runoff of residual ash and mobile ions like nitrate (NO_3_^-^), and lingering post-fire soil trace N gas emissions produced by microbial metabolism of post-fire soil N (e.g., nitrification and denitrification). Here, we hypothesize accelerated N trace gas emissions after wildfires from the ammonium (NH_4_^+^) left behind as ash that is nitrified to nitrite (NO_2_^-^) and NO_3_^-^ to also generate substrates for denitrification.

The extent to which soil physicochemical conditions and microbial processes shift after fire to influence soil trace N gas emissions may be influenced by burn severity (Goforth et al., 2005; Merino et al., 2018; Pressler et al., 2019; Smithwick et al., 2005; Ulery et al., 2017). Burn severity in soils, typically assessed by the loss of soil organic matter and ash deposition (Keeley, 2009), can determine the extent of soil pH increases, lowered C:N ratios (Dicen et al., 2020; Ulery et al., 2017), microbial biomass reduction (Caiafa et al., 2023; Holden & Treseder, 2013; Pulido-Chavez et al., 2022), and alter seed banks to affect plant community regeneration (Odion & Davis, 2000; Underwood et al., 2018). High severity fires could reduce microbial biomass enough to slow N cycling and reduce trace N gas emissions (Morishita et al., 2015), while low or medium severities could stimulate these processes by balancing increased soil N availability with microbial survival to maintain high N processing rates (Fierro & Castaldi, 2011; Gustine et al., 2022; Morishita et al., 2015; Strain et al., 2024). Because fires can have varied effects on soils depending on interactions with spatial variability in vegetation type, topography, fuel, and soil moisture (Evers et al., 2022; Keeley, 2009), gas fluxes may reflect fine-scale variability in soil burn severity. Thus, understanding soil burn severity interactions with soil gas emissions at the scale of a 1 m^2^ plot (representative of many field studies) could set expectations for variability and facilitate leveraging of existing coarse-scale assessments such as reported by the USDA Forest Service Burned Area Emergency Response (BAER) teams (typically at 30 m^2^ resolution; https://burnseverity.cr.usgs.gov; Parsons et al., 2010).

Over time after fire, soil N availability may decline as plants and microbes recover and take up soil N (Goodridge et al., 2018) and microbes undergo rapid succession (Pulido-Chavez et al., 2022), which may further alter nitrifier and denitrifier communities to moderate soil NO and N_2_O production. As post-fire ecosystem recovery progresses, early seral N-fixing plants may add to existing N pools with the potential to extend elevated N gas emissions (Erickson & Perakis, 2014) and further modify greenhouse gas balances (Kou-Giesbrecht et al., 2021). Thus, post-fire soil emissions of NO and N_2_O may change over time as plant and microbial succession progress, emphasizing the importance of understanding soil NO and N_2_O emissions over longer timescales post fire (previous studies in chaparral sampled over <1 year; Anderson et al., 1988; Anderson & Poth, 1989; Levine et al., 1988). To advance understanding of post-fire N cycling and drivers of soil NO and N_2_O emissions over time, we ask: i) how does wildfire influence soil emissions of NO and N_2_O over three years post fire? and ii) how do interactions between N emissions, soil burn severity, soil abiotic, and biotic factors drive ecosystem N losses post fire?

To answer these questions, we studied a chaparral shrubland adapted to wildfire and where NO and N_2_O emission pulses are commonly observed after wetting dry soils. We measured fluxes of NO and N_2_O in the laboratory after wetting soils collected across a burn severity gradient assessed at the 1 m^2^ scale after the 2018 Holy Fire burned in the Cleveland National Forest in Southern California. We modeled the impacts of a suite of fire-impacted variables on trace N gas emissions to better understand which factors drive post-fire soil emissions of NO and N_2_O in chaparral. We hypothesized that: i) soil emissions of NO and N_2_O would increase post fire, decreasing over time as N substrates are exhausted, ii) soil emissions of NO and N_2_O would be highest in soils that burned at medium severity due to the combination of high pH and NH_4_^+^ availability with higher microbial survival, and iii) NO and N_2_O emissions would be driven by pH, NH_4_^+^, NO_3_^-^, and microbial biomass.

## 2. Materials and Methods

### 2.1 Field Site Description

From August 6th to September 13th, 2018, the Holy Fire burned 94 km^2^ of the Cleveland National Forest in Southern California. Two weeks after the fire was extinguished, a network of nine plots, six burned and three unburned, each with four 1-m^2^ subplots (n = 36) located 5 m from the plot center in each cardinal direction (**Figure S1**) were established in manzanita- dominated stands (*Arctostaphylos glandulosa*) on south-facing slopes (slope 33 – 57%) at an average elevation of 1,200 m as previously published (Pulido-Chavez et al. 2022). Climate is Mediterranean with the dry season occurring May through November with mean annual temperature 17.5°C ± 0.68°C. The mean annual precipitation over the study period was 320 mm (El Cariso weather station; <u>raws.dri.edu</u>). Atmospheric N deposition at this site is estimated to fall between 8−12 kg N ha^-1^ y^-1^ (Schwede & Lear, 2014). Soils were classified as Entisols (Typic Xerorthents) mapped in the Cieneba series and Mollisols (Lithic Haploxerolls) mapped in the Friant Series (**Table S1;** Soil Survey Staff, 2024). Because the extent of the area burned limited access to nearby unburned manzanita-dominated stands, it was not possible to establish unburned plots in the Entisols. However, when Entisols were removed from the dataset, fire effects were similar for the majority of soil variables (**Table S2**) implying that the effect of fire overwhelmed differences between soil series. Thus, we include both soil types in our analyses.

### 2.2 In situ soil sampling and soil and vegetation monitoring

Burned plots were reported as moderate soil burn severity by the USDA FS BAER team (at 30-m^2^ resolution; Nicita & Halverson, 2018). To capture heterogeneity at the 1-m^2^ subplot scale, we measured ash depth (cm), averaged over three measurements per subplot, two weeks post fire as a proxy of soil burn severity as previously published (Pulido-Chavez et al., 2022). To approximate BAER categories at the 1-m^2^ scale, we also binned ash depths into categories of Low (0.01 – 1.3 cm), Medium (1.4 – 3.0 cm), and High (3.1 – 12.0 cm; **Table S3**) burn severity as outlined in the USDA FS Field Guide for Mapping Post-Fire Soil Burn Severity (Parsons et al., 2010).

We analyzed soil samples collected at 10 timepoints including 4 mo, 6 mo, 9 mo, 1 y, 1 y 4 mo, 1 y 8 mo, 2 y, 2 y 4 mo, 2 y 7 mo, and 3 y post fire. Soil cores were taken from the top 10 cm of mineral soil (A horizon; duff and ash were removed if present) using a 7.5 cm diameter auger and wiped with 70% ethanol between subplots. Soil cores were transported to the University of California Riverside (UCR) on ice and sieved (2 mm) within 24 h. Subsamples for qPCR and Illumina MiSeq sequencing were immediately stored at -80 °C for DNA extraction (see supplementary **Methods S1**) and separate subsamples immediately oven-dried to measure field soil gravimetric water content (GWC; ∼10 g). Freshly collected soil samples were analyzed for GWC, soil extractable NH_4_^+^, nitrite (NO_2_^−^), NO_3_^−^ (within 24 h), and total soil microbial biomass C and N (within 72 h) after which soils were stored by air-drying at 25 °C.

Soil extractable NH_4_ ^+^ and NO_3_^-^ concentrations were measured by extraction with 2 M KCl (Maynard et al., 2008). Water extractable NO_2_^-^ was extracted with nanopure water because KCl underestimates nitrite concentrations (Homyak et al., 2015). We also measured *in situ* net N transformation rates at two subplots at each plot (N and S; n = 18) at 1 y 4 mo, 1 y 8 mo, and 2 y post fire by homogenizing 10-cm × 7.5-cm soil cores and burying half in a sealed polyethylene bag in the original core location, while the other half was transported to UCR in a cooler to measure the initial concentration of NH_4_ ^+^ and NO_3_^-^ within 24 h (Eno, 1960). Buried bags were removed after approximately 1 month and re-measured for extractable NH_4_ ^+^ and NO_3_^−^. *In situ* net N transformation rates were calculated as the difference in NH_4_^+^ (net NH_4_^+^ production rate) or NO_3_^−^ (net nitrification rate) between final and initial measurements over 30 days. Soil extracts were analyzed colorimetrically at the Environmental Sciences Research Laboratory (ESRL; https://envisci.ucr.edu/research/environmental-sciences-research-laboratory-esrl) at UCR for NH_4_^+^ (SEAL method Environmental Protection Agency (EPA)-126-A; detection limit 0.05 mg/L), NO_3_^−^ (SEAL method EPA-129-A; detection limit 0.03 mg/L), and NO_2_^-^ (SEAL method EPA-137-A; 0.0001 mg/L) using a SEAL AQ-2 discrete analyzer.

Soil microbial biomass C (MBC) and microbial biomass N (MBN) were measured using a chloroform slurry-extraction method in 0.5 M K_2_SO_4_ (Fierer & Schimel, 2002) where extractable C and N are measured on a total organic C and N analyzer (Shimadzu Corporation, Series V Model CSN) and the contents of CHCl_3_-fumigated extracts are compared to unfumigated extracts. We did not correct MBC or MBN for extraction efficiency (Vance et al., 1987) and thus our measurements represent a flush of C and N after fumigation.

Bulk soil % C, % N, δ^13^C, and δ^15^N were measured with an elemental analyzer (Costech) coupled with an isotope ratio mass spectrometer (Thermo Delta V, Thermo Fisher Scientific, Woltham, MA) at the UC Riverside Facility for Isotope Ratio Mass Spectrometry (FIRMS; https://ccb.ucr.edu/facilities/firms). Water holding capacity (WHC) was calculated as the amount of water a saturated soil could hold after freely draining in a sealed container to prevent evaporation for 48 h. Soil texture (% sand, silt, and clay) was measured using the hydrometer method (Gavlak et al., 2005) and bulk density was measured by weighing oven-dried intact soil cores of a known volume. Soil pH was measured in a 1:2 solution of soil in deionized water with a pH meter (Thermo Scientific; Orion Versastar Pro meter equipped with ROSS Ultra pH electrode).

Vegetation species composition was monitored over the study period as the absolute percent cover of each plant species present at designated 1 m^2^ plots adjacent to soil sampling plots (**Figure S1B**). We selected measurements at 1 y, 2 y, and 1 y 8 mo post fire, which approximately matched those selected for soil gas flux analysis. To estimate the effect of N-fixing plants on N cycling, plant species that are known to form associations with N-fixing bacteria were identified within this data set: *Acmispon glaber* (Fabaceae)*, Acmispon strigosus, Ceanothus crassifolius* (Rhamnaceae)*, Ceanothus leucodermis,* and *Ceanothus oliganthus*. The percent composition of N-fixing species were summed to represent total percent cover of N-fixing plants at each plot.

### 2.3 Laboratory measurements of NO and N_2_O and extractable N

Soils were selected for NO and N_2_O flux measurements (n = 24 burned, n = 6 unburned) from air-dried samples collected in September at one year, two years, and three years post fire to represent conditions at the end of the dry season (hereafter referred to as the end of years 1−3 post fire). We focused on the dry season to better understand the potential for burned soils to stimulate N emissions during the dry-to-wet seasonal transition, a period well recognized to stimulate soil N emission pulses in drylands (Homyak et al., 2016, 2017; Krichels et al., 2022, 2024). To simulate the first end-of-dry-season rain event and activate microbial processes (Homyak et al., 2017), we wetted 50 g of air-dried soil to 75% WHC in 120-mL Mason jars and immediately measured soil fluxes of NO and N_2_O for 48 h by multiplexing to a continuous flow trace gas analyzer system as described in detail by Krichels et al. (2024). This consists of a recirculating sample loop controlled by a multiplexer (LI-8150, LI-COR Biosciences) connected to a laser isotopic N_2_O analyzer (Los Gatos Research, Inc.; Model 914–0027), an infrared CO_2_/H_2_O analyzer (LI-8100, LI-COR Biosciences), and a chemiluminescent NO_2_ analyzer (Scintrex LMA-3D, Unisearch Associates, Canada) fitted with a CrO_3_ converter to oxidize NO to NO_2_. Full flux measurements are described in detail in supplementary materials (**Methods S2**).

To avoid including the less stable periods of flux ramp up and decline, flux data were trimmed to 40 h by removing the first and last 4 h of measurement time for both NO and N_2_O before calculating cumulative emissions via trapezoidal integration (trapZ function in R; R Core Team, 2024). Water content was tracked by weighing every 24 h. Soil NH_4_^+^ and NO_3_^-^ extractions were performed pre- and post-48-h incubations parallel to flux measurements to calculate net N transformation rates as the change between final and initial measurements in soil NH_4_^+^ (net NH_4_^+^ production rate) or soil NO_3_^−^ (net nitrification rate) per day.

### 2.4 Natural abundance isotopic composition of N_2_O

To assess possible sources of N_2_O emissions, we measured isotopocules of N_2_O (δ^15^N_2_O_bulk_, δN_2_^18^O_bulk_, and site preference (SP) δ^15^N_2_O_SP_, which reflects the placement of ^15^N in the central (α) and peripheral (β) positions in the N_2_O molecule; where δ = (((Rsample/Rstandard)−1)×1000) in units of ‰ and R = ^15^N/^14^N or ^18^O/^16^O) from six soils chosen to represent the two highest cumulative N_2_O fluxes in each year and one unburned soil from each year (soils from year 2 did not produce enough N_2_O for isotope analysis; n = 5 burned, 2 unburned). Briefly, 50g of each soil was incubated at 75% WHC in the lab and headspace was captured in a 1-L gas-tight bag to measure N_2_O on a cavity ringdown infrared N_2_O analyzer (Los Gatos Research, Inc.). Methods are described in full in supplementary materials (**Methods S3**).

Sample isotopic values were compared to previously published ranges of isotopomers of N_2_O (Lewicki et al., 2022; Yu et al., 2020) for distinct microbial processes such as nitrification (Ni), nitrifier denitrification (nD), bacterial denitrification (bD), and fungal denitrification (fD). We then used the publicly available FRAME model to partition fractional contributions of N_2_O production processes (Ni, fD, nD, bD) and the residual N_2_O fraction (r) to estimate the proportion of N_2_O reduction (Lewicki et al., 2022; Well & Flessa, 2010; Yu et al., 2020).

### 2.5 Statistical analyses

All statistical analyses were performed in R 4.3.1 (R core team). Linear mixed effects models (LME; lme4 package in R; Bates et al. 2015) were applied in three different analyses: **1)** to test for differences between variables across burned and unburned plots over sampling times at the end years 1−3 (**Table 1**) and at individual sampling time points over a finer temporal sampling scheme using post-hoc analysis (**Figures 2, 3, 4, & 5**); **2)** to determine differences across soil burn severity categories compared to unburned (**Figure 2**); and **3)** to understand the relative importances of factors affecting fluxes of NO and N_2_O using a multiple regression approach (**Figure 6**). All models used restricted estimation maximum likelihood (REML; lme4 package in R) and included subplot as a random effect (models that included plot and subplot were not different from models that only included subplot), a first-order autoregressive correlation structure to account for repeated sampling over time (Zuur et al., 2009, https://m-clark.github.io/mixed-models-with-R/), and a weighting structure to account for variance by soil type. For analysis **1**, significant differences between variables in burned and unburned plots sampled at the end of years 1−3 were assessed with a categorical factor for burn treatment and an interaction with time. If there was a significant interaction of burn and time (p ≤ 0.05), then to test for individual differences between burned and unburned soils in a specific sampling time, post-hoc analysis was performed using the “emmeans” package in R (Lenth et al., 2023). For analysis **2**, significant differences between unburned and each burn severity treatment were assessed by modeling each variable with a categorical factor for soil burn severity (low, medium, and high) across all three years of the study. The effects of burning over time and across different burn severities were modeled separately for fluxes of NO and N_2_O with a log transformation of each response variable to meet normality assumptions.

**Figure 2.**
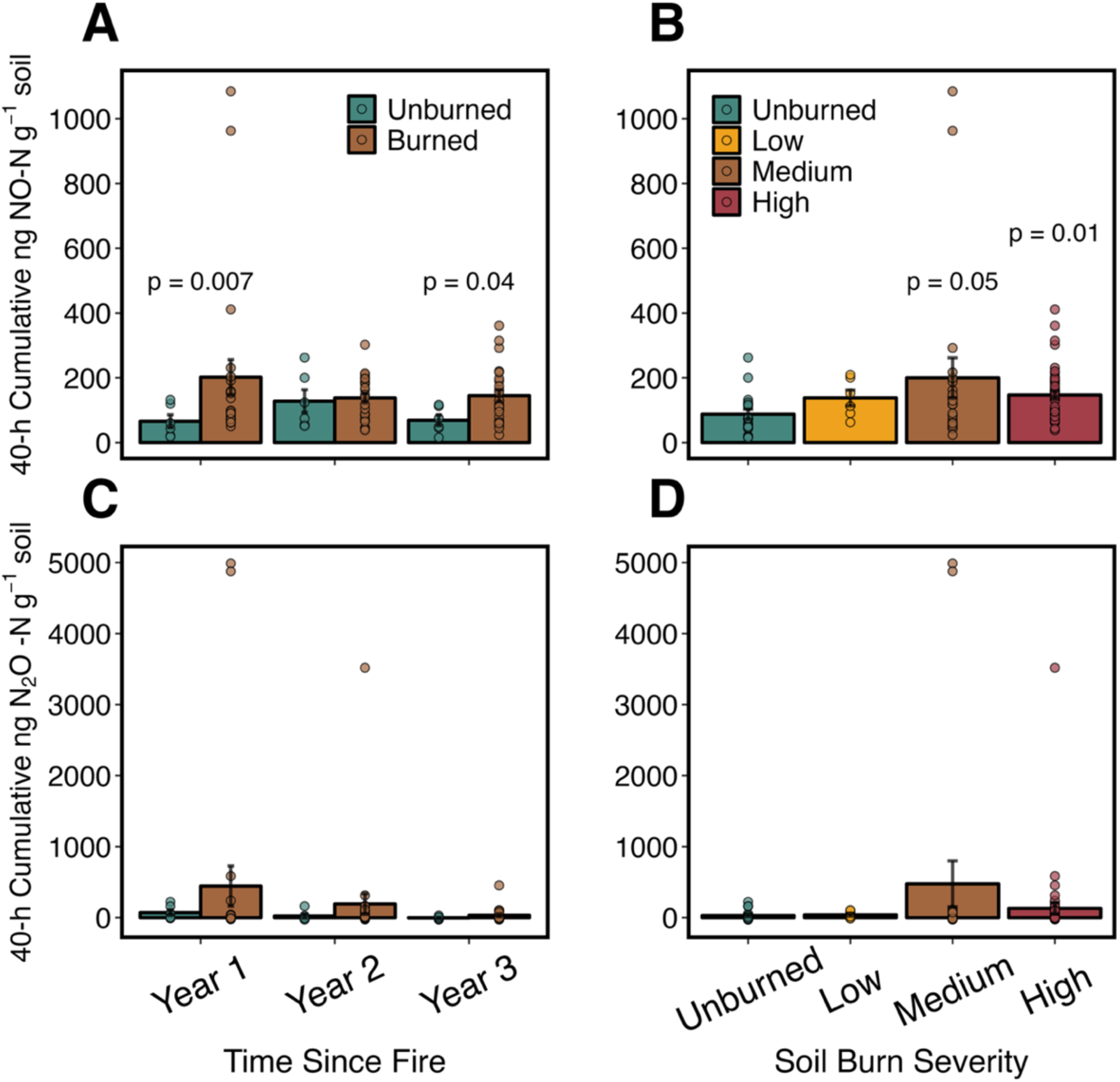
Cumulative soil NO and N_2_O emissions measured over 40-h lab incubations. Linear mixed effects models that accounted for soil series, plot, and repeated sampling over time were used to estimate the effects of burning (**A, C**) or burn severity (**B, D**) on fluxes of NO and N_2_O compared to unburned. P-values are post-hoc comparisons performed when there was significant interaction between burning and time for **A** or **C**, to show the significance of the effects of burning compared to unburned for individual years (full interactions and estimates in **Table 1**). Dots represent measurements from individual soils. Error bars represent standard errors, for each timepoint n = 6 unburned, n = 24 burned.

**Table 1.** Summary of variable means in burned and unburned plots combining all data from the end of years one, two, and three post-fire. Distinction is made between variables measured on field-fresh soil (*in situ*) and in laboratory incubations. Linear mixed effects models that accounted for soil series, plot, and repeated sampling over time were used to test for differences between burned/unburned groups and interactions over time. “Estimate” is the modeled average unit change in burned compared to unburned plots over all three years combined, while p-values show significant differences between burned and unburned groups (“Burn”), over time (“Time”), and the interaction of burning and time (“Burn ξ Time”). Bulk density was measured at only one sampling time one-month post-fire, so no time interactions are shown.

| Table 1 |  | Burned |  |  | Unburned |  |  | Estimate | P-values |  |  |
| --- | --- | --- | --- | --- | --- | --- | --- | --- | --- | --- | --- |
| Variable (*in situ*) | unit | Mean | SE | n | Mean | SE | n | Burn | Burn | Time | Burn × Time |
| Bulk density | g cm<sup>-3</sup> | 0.82 | 0.03 | 24 | 0.56 | 0.05 | 12 | 0.27 | <0.001 | NA | NA |
| Soil Moisture | g water g soil<sup>-1</sup> | 0.020 | 0.002 | 72 | 0.052 | 0.006 | 36 | -0.036 | <0.001 | 0.162 | 0.228 |
| pH |  | 6.58 | 0.06 | 72 | 6.13 | 0.05 | 35 | 0.43 | <0.001 | 0.326 | 0.078 |
| Bulk soil C | mg C g soil<sup>-1</sup> | 27.34 | 1.63 | 72 | 46.26 | 7.26 | 18 | -22.06 | 0.007 | 0.225 | 0.001 |
| Bulk soil N | mg N g soil<sup>-1</sup> | 1.45 | 0.09 | 72 | 1.87 | 0.26 | 18 | -0.44 | 0.183 | 0.046 | 0.003 |
| C:N | ratio | 22.4 | 0.5 | 72 | 27.8 | 1.2 | 18 | -5.0 | 0.002 | <0.001 | 0.455 |
| Bulk soil $\delta^{15}\text{N}$ | ‰ | 2.10 | 0.13 | 72 | 1.80 | 0.38 | 18 | 0.24 | 0.508 | 0.019 | 0.008 |
| Bulk soil $\delta^{13}\text{C}$ | ‰ | -26.63 | 0.10 | 72 | -26.87 | 0.35 | 18 | 0.14 | 0.403 | <0.001 | <0.001 |
| Ammonium | μg NH<sub>4</sub><sup>+</sup>-N g soil<sup>-1</sup> | 16.1 | 1.7 | 72 | 5.8 | 2.5 | 36 | 8.8 | 0.012 | <0.001 | 0.218 |
| Nitrate | μg NO<sub>3</sub><sup>-</sup>-N g soil<sup>-1</sup> | 9.1 | 1.7 | 72 | 0.7 | 0.2 | 36 | 7.5 | 0.002 | <0.001 | 0.003 |
| Microbial biomass C | μg C g soil<sup>-1</sup> | 135.1 | 9.9 | 72 | 370.8 | 50.3 | 36 | -238.8 | <0.001 | <0.001 | 0.031 |
| Microbial biomass N | μg N g soil<sup>-1</sup> | 5.1 | 0.7 | 72 | 21.9 | 3.5 | 36 | -17.4 | <0.001 | 0.001 | 0.192 |
| Bacterial abundance | gene copy number | 1.74E+08 | 1.02E+07 | 70 | 5.57E+08 | 2.69E+08 | 35 | -4.28E+08 | 0.039 | 0.346 | 0.030 |
| Fungal abundance | gene copy number | 1.54E+07 | 1.81E+06 | 70 | 7.80E+07 | 8.61E+06 | 35 | -6.04E+07 | <0.001 | 0.030 | 0.168 |
| Bacterial Richness | Total no. bacterial sp. | 547 | 31 | 70 | 575 | 36 | 31 | -30 | 0.566 | <0.001 | 0.433 |
| Fungal Richness | Total no. fungal sp. | 103 | 5 | 70 | 286 | 15 | 35 | -182 | <0.001 | 0.345 | 0.184 |
| N-fixing plants | % cover N fixing sp. | 5.7 | 1.5 | 72 | 3.2 | 1.0 | 36 | 2.6 | 0.375 | <0.001 | 0.084 |
| Vegetation Richness | Total no. plant species | 4 | 0.3 | 72 | 2 | 0.2 | 18 | 1.5 | 0.064 | <0.001 | 0.145 |
| Variable (Laboratory) |  | Mean | SE | n | Mean | SE | n | Burn | Burn | Time | Burn × Time |
| CO<sub>2</sub> flux | ng CO<sub>2</sub>-C g<sup>-1</sup>soil | 153809 | 11156 | 72 | 170621 | 23694 | 18 | -33052 | 0.523 | 0.683 | 0.180 |
| NO flux | ng NO-N g<sup>-1</sup>soil | 161.4 | 19.6 | 72 | 87.4 | 15.4 | 18 | 79.7 | 0.007 | 0.621 | 0.026 |
| N<sub>2</sub>O flux | ng N<sub>2</sub>O-N g<sup>-1</sup>soil | 224.6 | 106.8 | 72 | 30.9 | 17.4 | 18 | 195.6 | 0.458 | 0.136 | 0.376 |
| NO:N<sub>2</sub>O | ratio | 17.0 | 19.1 | 72 | -0.1 | 2.1 | 18 | 4.1 | 0.922 | 0.217 | 0.937 |
| Net mineralization | μg N g soil<sup>-1</sup> day<sup>-1</sup> | 4.67 | 1.88 | 70 | 3.55 | 1.03 | 17 | 1.09 | 0.667 | 0.002 | 0.174 |
| Net nitrification | μg NO<sub>3</sub><sup>-</sup>-N g soil<sup>-1</sup> day<sup>-1</sup> | 4.08 | 1.74 | 70 | 0.25 | 0.19 | 17 | 2.99 | 0.328 | 0.008 | 0.350 |
| Net NH<sub>4</sub><sup>+</sup> production | μg NH<sub>4</sub><sup>+</sup>-N g soil<sup>-1</sup> day<sup>-1</sup> | 0.59 | 0.90 | 70 | 3.30 | 1.02 | 17 | -2.45 | 0.176 | 0.036 | 0.668 |

To estimate the relative importance of factors driving NO and N_2_O emissions (analysis **3**), we constructed global linear mixed effects models for each response variable (NO or N_2_O) by selecting a suite of predictor variables that best represented potential controls on emissions.

Pearson’s correlation test was used to eliminate colinear variables (for example, percent clay was chosen to represent texture due to collinearity; **Figure S2**) and variables from similar informational categories were further narrowed down based on individual correlations (for example, bacterial and fungal abundance were chosen over richness due to better correlation and lack of significance when richness was included in global models) or better representation of bulk processes (for example, net N transformation rates were chosen over initial NH_4_^+^ and NO_3_^-^ concentrations). Final global models for each gas (NO, N_2_O) contained ten predictor variables (pH, GWC, C:N, MBC, net NH_4_^+^ production rate, net nitrification rate, % cover of N-fixing species, % clay content, bacterial abundance, and fungal abundance; **Eq. S1, S2**) and log-transformations were applied to NO and N_2_O fluxes to fit the assumption of normal distributions. To leverage the entire dataset and understand bulk drivers of NO and N_2_O across a range of burn severities, we considered burning as a gradient from unburned to low, medium, and high severity and included all plots in the analysis. Global models were assessed for fit using quantile-quantile plots and histograms of model residuals. Model selection was performed using a maximum likelihood approach (“dredge()” function in the MuMIn package; Bartoń 2010). Final models were selected using Aikake Information Criterion (AIC) and similar models (delta <2) were averaged (**Table S4;** Symonds & Moussalli, 2011). To allow direct comparison of coefficients for visualization (**Figure 6**), we scaled the dependent variables by subtracting the mean and dividing by the standard deviation.

## 3. Results

### 3.1 Laboratory fluxes of soil NO and N_2_O

Wildfire significantly increased soil NO emissions when averaged across all years (**Figure 2A**; burn: p = 0.007) and the effect varied as a function of time (burn × time: p = 0.03; **Table 1**) such that soil NO fluxes were significantly higher in burned soils in years one (p = 0.007) and three (p = 0.04) compared to unburned, with no difference between burned and unburned plots in year two (**Figure 2A**). Cumulative 40-h NO fluxes over the three-year period were roughly two times higher in burned plots (hereafter, average ± standard error; 161 ± 19 ng NO-N g^-1^ soil) relative to unburned (88 ± 15 ng NO-N g^-1^ soil). Further, soils that burned at Medium (p = 0.04) and High (p = 0.01) severity had significantly higher NO emissions compared to unburned plots, whereas soils that burned at Low severity did not differ significantly from unburned plots (p = 0.7; **Figure 2B; Table S3**). The highest soil flux recorded for NO reached 1,084 ng NO-N g^-1^ soil (measured in year one, at Medium severity).

Due to high variation in soil N_2_O emissions, we did not detect a significant effect of wildfire (**Figure 2C**; average 40-h cumulative fluxes over all years were 224 ± 106 ng N_2_O-N g^-1^ soil in burned plots and 30 ± 17 ng N_2_O-N g^-1^ soil in unburned plots; **Table 1**). However, only burned soils produced on occasion very high N_2_O emissions that were identified as statistical outliers by multiple tests (Interquartile Range, Percentiles of 97.5 and 2.5, and z-scores >3) and were up to 165 × higher than the average in unburned plots (> 3,500 ng N_2_O-N g^-1^ soil in burned plots relative to < 100 ng N_2_O-N g^-1^ soil in unburned plots; **Figure 2C&D**). NO:N_2_O ratios averaged -0.1 ± 2.1 in unburned plots and 17 ± 19.1 in burned plots but did not differ significantly (**Table 1**).

### 3.2 Soil N Cycling

*In situ* soil extractable NH_4_^+^ and NO_3_^-^ concentrations increased post fire (**Figure 3**) and then declined to match unburned soils by 1 y 8 mo for NH_4_^+^ (**Figure 3A)** and by 1y 4 mo for NO_3_^-^ (**Figure 3B**). When all soils measured at the end of years 1–3 post fire were compared to unburned, soil extractable NH_4_^+^ was elevated by an average of 10 µg NH_4_^+^-N g soil^-1^ in burned plots (**Table 1**; burn: p = 0.01) with the highest average falling in the High burn severity category (**Table S3**), which was the only burn severity category that differed significantly from unburned (**Figure S3A**; p = 0.02). The peak value (230 µg NH_4_^+^-N g soil^-1^) for an individual soil core was measured in a plot that burned at high severity four months after fire. When seasonal sampling times were considered, the highest concentrations of soil extractable NH_4_^+^ measured during year one fell at 4 mo and 9 mo post fire in burned plots (**Figure 3A**). Similar to soil extractable NH_4_^+^, soil extractable NO_3_^-^ averaged 8 µg NO_3_^-^-N g soil^-1^ higher in burned plots when all soils measured at the end of years 1–3 post fire were compared to unburned (**Table 1**; burn: p = 0.002; burn × time: p = 0.003), and the highest averages in the Medium and High burn severity categories (**Table S3**) with only the High burn severity category differing significantly from unburned (**Figure S3B**; p = 0.02). However, we found that soil extractable NO_3_^-^ in burned plots varied strongly over time and did not differ from unburned for the first 6 mo post fire but reached a peak average of 17 times higher than soils from unburned plots at 9 mo (**Figure 3B**). The highest individual measurement for soil extractable NO_3_^-^ was measured in a plot that burned at high severity in the same 9 mo sampling time (138 µg NO_3_^-^-N g soil^-1^).

**Figure 3.**
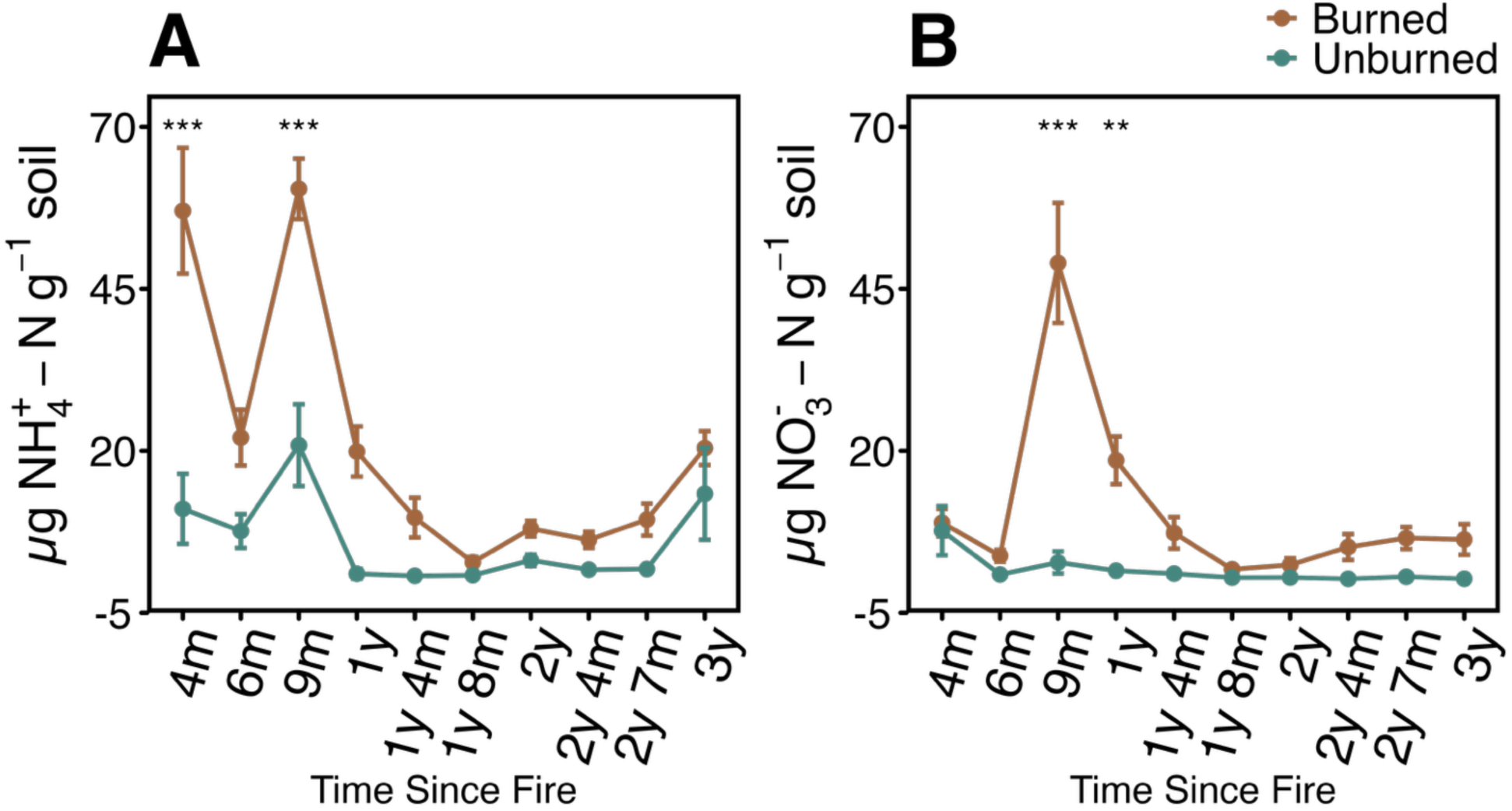
Post-fire soil N pools measured in situ from burned and unburned plots over the 3-year study period: soil extractable ammonium (NH_4_^+^; **A**), and extractable nitrate (NO_3_^-^; **B**). Points represent means for each timepoint, error bars are standard error, and for each timepoint n = 12 unburned, n = 24 burned. Linear mixed effects models were used to account for repeated sampling over time and different variances between soil series and plots to estimate the effects of burning on each variable compared to unburned (**Table 1**). Post-hoc comparisons show the significance of the effects of burning compared to unburned at each sampling time with asterisk signifying p-values of: ‘***’ <0.001, ‘**’ <0.01, ‘*’ <0.05. The wildfire was extinguished on September 13, 2018.

*In situ* net N transformation rates measured during year two post fire were more negative in burned plots compared to unburned for net NH_4_^+^ production (**Figure 4A**: burn: p = 0.02), indicating gross N consumption exceeded gross N production. Net nitrification, however, did not differ between burned and unburned in bulk or over time (**Figure 4B**: burn: p = 0.4). Net N transformations measured in the laboratory over 48 h concurrently with NO and N_2_O emissions did not differ between burned and unburned plots (**Table 1**, **Figure 4C&D**).

**Figure 4.**
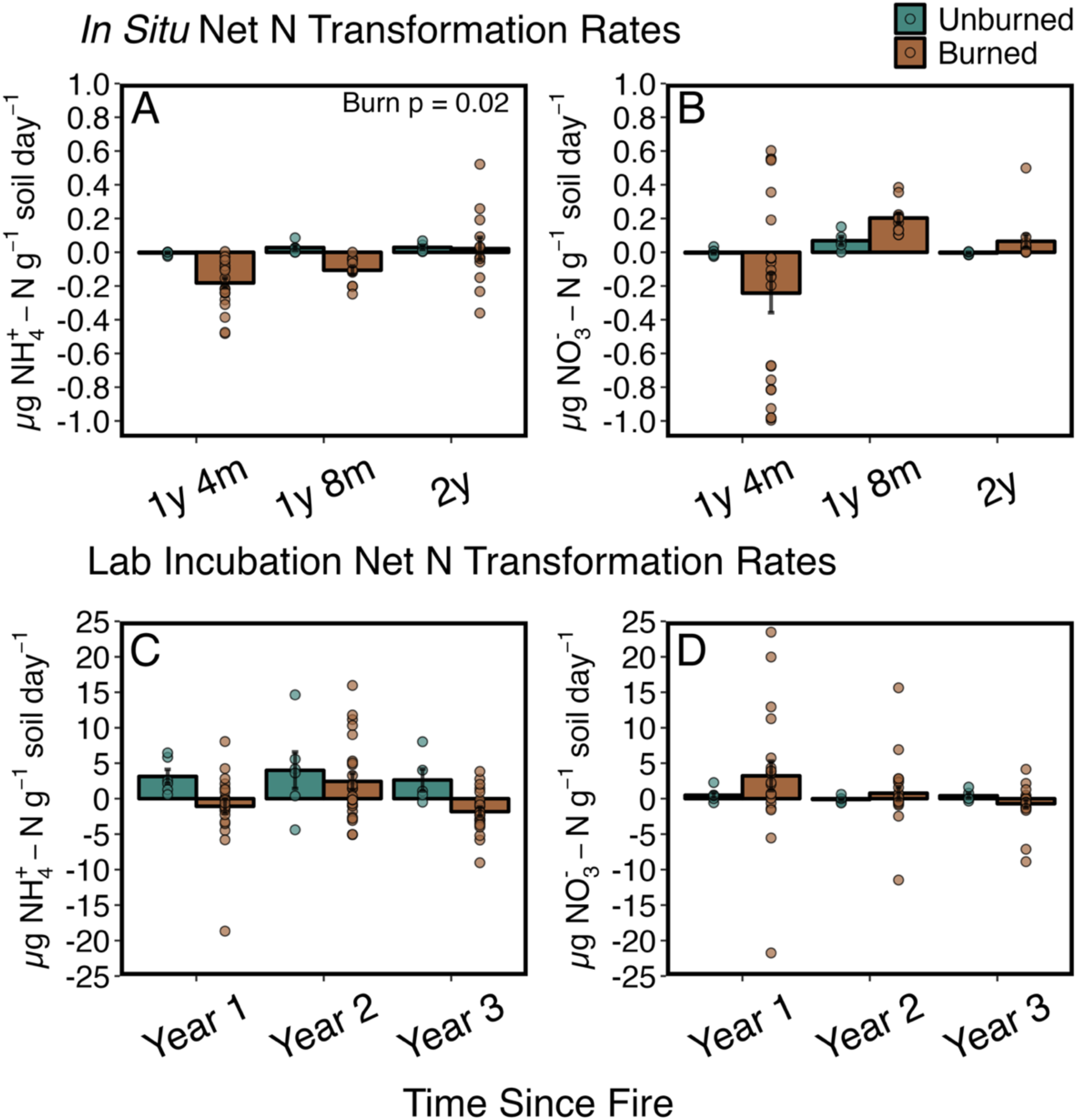
Net NH_4_^+^ production and net nitrification rates measured *in situ* over one-month periods using buried soil cores in the field during the second year post fire (**A, B**), and during 48-h soil laboratory incubations from soil samples collected at the end of years 1−3 post fire (**C, D**). P-value in top right **A** shows significance of burning alone (“Burn”). Bars represent mean and error bars represent standard errors. For **A** and **B**: for each timepoint: n = 24 burned, n = 12 unburned; and for **C** and **D**: for each timepoint n = 24 burned, n = 6 unburned.

### 3.3 Soil biogeochemical properties

Soil bulk density was significantly higher in burned soils (**Table 1**, burn: p = 0.01). *In situ* soil gravimetric water content (GWC) declined significantly by an average of 0.03 g water g^-1^ soil in burned plots measured at the end of years 1–3 post fire (**Table 1**; burn: p < 0.001). When all sampling times were considered, decreases in GWC in burned plots compared to unburned were strongest during sampling times that corresponded to months with the highest total precipitation (**Figure 5A&B**). Soil pH increased significantly after fire with an average 0.5 unit increase in burned plots over three years (**Table 1**; burn: p < 0.001) and remained significantly elevated at every sampling time point (**Figure 5C**) with the strongest increases in Medium and High burn severity categories (**Figure S3C**). Bulk soil total C decreased significantly by an average of 19 mg C g soil^-1^ in burned soils (**Table 1**; burn: p = 0.007), but bulk soil N only differed significantly from unburned in year two, with a decrease in burned plots by an average of 1.1 mg N g soil^-1^ (**Table 1**; burn × time: p = 0.003; post-hoc year 2: p = 0.007). Bulk soil total carbon to nitrogen (C:N) ratios were significantly lower in burned plots compared to unburned (**Figure S4A**) with an average 25% decrease over all three years (**Table 1**; burn: p = 0.002).

**Figure 5.**
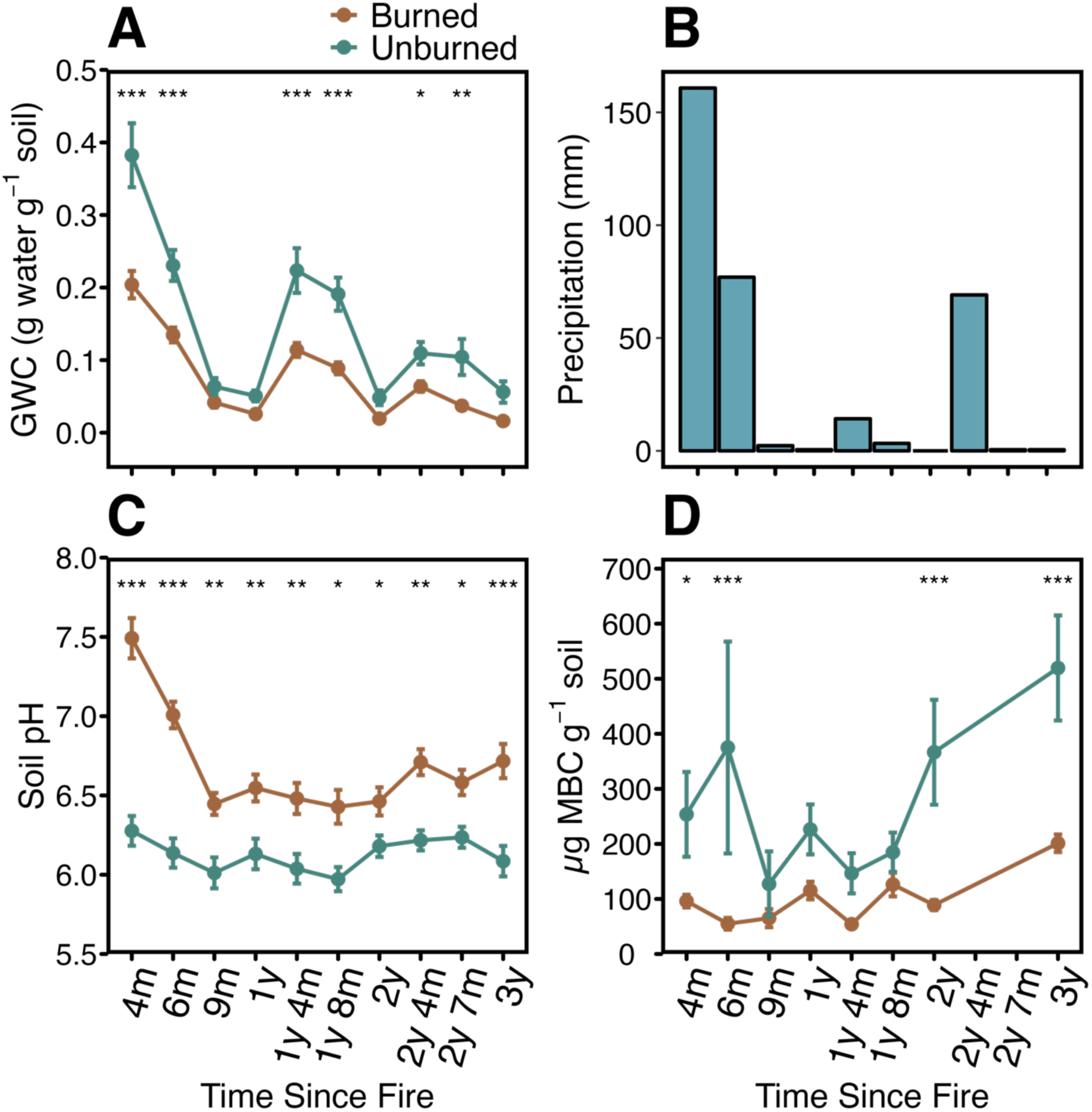
Post-fire monitoring of *in situ* gravimetric water content (GWC; **A**), the total monthly precipitation (mm) that fell in the month of each sampling time point (**B**), soil pH (**C**), and *in situ* microbial biomass C (MBC; **D**), measured on soils from burned and unburned plots over the 3-year study period. Points represent means, error bars are standard error, and for each timepoint n = 12 unburned, n = 24 burned. Post-hoc comparisons show the significance of the effects of burning compared to unburned at each sampling timepoint with asterisk signifying p-values of: ‘***’ <0.001, ‘**’ <0.01, ‘*’ <0.05. The wildfire was extinguished on September 13, 2018.

**Figure 6.**
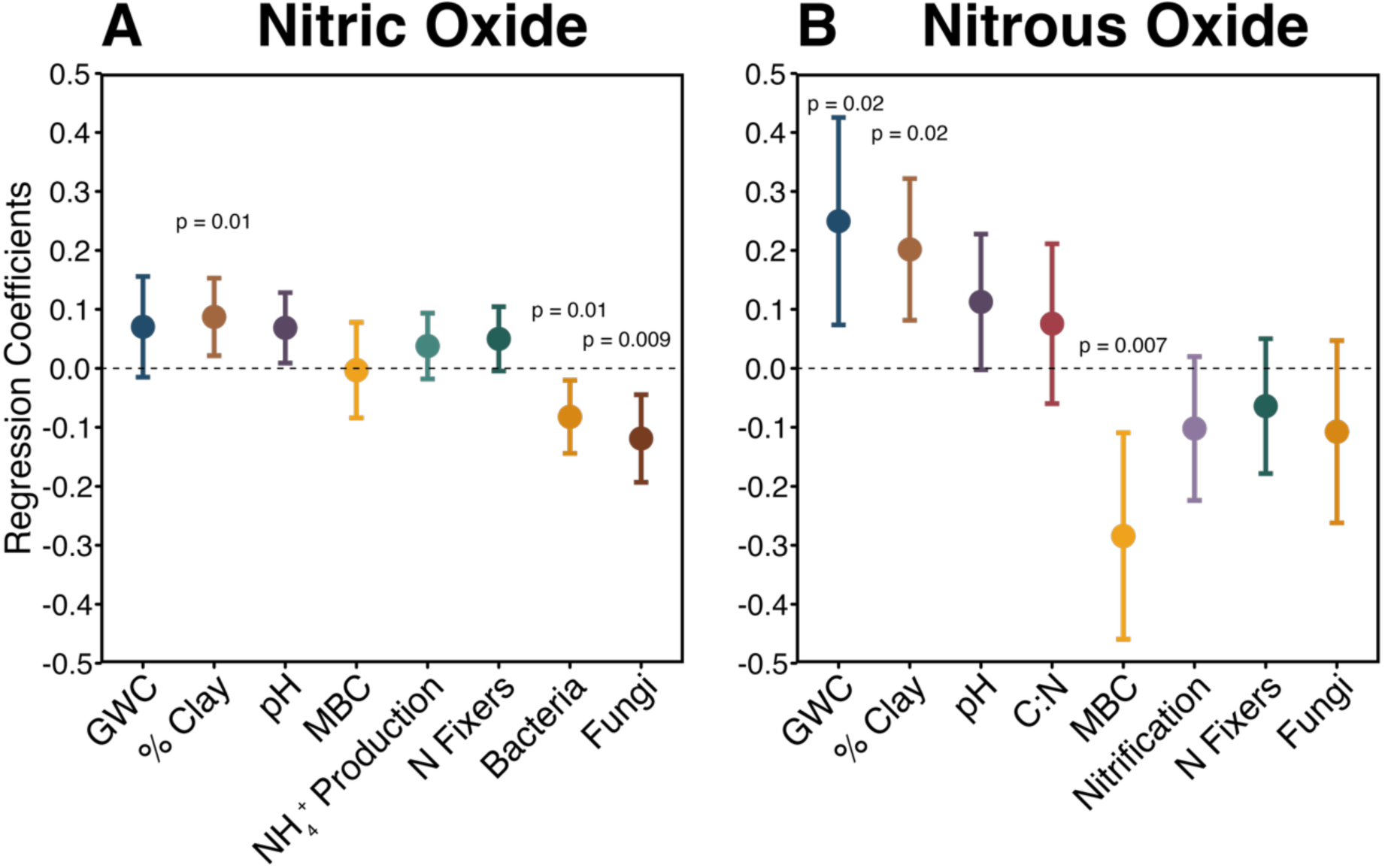
Summary of regression coefficients for abiotic and biotic variables that mediate cumulative soil emissions of NO (**A**) and N_2_O (**B**). Fluxes were modeled separately using a linear mixed effects multiple regression approach that accounted for plot, soil type, and repeated measures. For visualization, data were scaled by subtracting the mean and dividing by the standard deviation to standardize the estimated regression coefficients so that the distance each estimate is from 0 ranks the size of the effect on the response (full model averages in **Table S4**). P-values show significance of averaged modeled estimates. Variables measured in tandem with gases over the course of the 48-h incubation included gravimetric water content (g water g^-1^ soil; “**GWC**”), “**NH_4_^+^ production**” (μg NH_4_^+^-N g^-1^ soil day^-1^), and “**Nitrification**” (μg NO_3_^-^-N g^-1^ soil day^-1^). *In situ* variables included “**N fixers**” (percent cover of N-fixing plant species), “**Bacteria**” and “**Fungi**” (bacterial and fungal abundance measured as gene copy number), and microbial biomass C (µg C g soil^-1^; “**MBC**”). Soil pH, % clay, and C:N ratio were measured on air-dried soils. Error bars represent 95% confidence intervals of the regression coefficients.

Bulk soil δ^13^C and δ^15^N changed in response to fire and time, with bulk soil δ^15^N becoming significantly enriched in burned plots in year two (**Figure S4B**; burn × time: p = 0.008; post-hoc for year two: p = 0.01), and bulk soil δ^13^C becoming significantly more enriched in burned plots in year one (**Figure S4C**; burn × time: p < 0.001; post-hoc for year one: p = 0.002).

### 3.4 Soil microbial and vegetation monitoring

Burning significantly decreased bulk soil microbial biomass C (MBC) by an average of 235 µg C g soil^-1^ measured at the end of years 1–3 post fire (**Table 1**; burn: p < 0.001; burn × time: p = 0.03) and microbial biomass N by an average of 17 µg N g soil^-1^ (**Table 1**; burn: p < 0.001). MBC was significantly lower in the burned plots compared to unburned up to 6 mo post fire, did not differ from unburned from 9 mo to 1y 8 mo post fire, but significantly diverged from unburned again in year two and showed little evidence of recovery by year three (**Figure 5D**).

MBC differed from unburned across all burn severity categories, but did not differ between Low, Medium, and High (**Figure S3D**). Bacterial abundances were significantly lower in burned plots in years 1 and 3 (**Figure S5A**; **Table 1**; burn p = 0.04) and differed from unburned at the Low and High burn severity categories (**Figure S5B)**. Fungal abundance was lower in burned plots at every sampling time (**Figure S5C**; **Table 1**; burn: p < 0.001) and was significantly lower than unburned at all burn severity categories (**Figure S5D**). Bacterial richness did not differ significantly between burned and unburned at the end of years 1–3 post fire (**Table 1**; burn p = 0.5); however, fungal richness was lower in burned plots in all years (**Table 1**; burn: p < 0.001). Although the interaction between burn and time was only significant at p = 0.08 for the summed percent cover of N-fixing plant species (**Table 1**), N-fixing plant cover increased in burned plots over the three years (**Figure S6**; time: p < 0.001).

### 3.5 Linear Mixed Effects Models for NO and N_2_O

Using a linear mixed effects approach to understand the relative importance of ten potential explanatory variables (pH, GWC, C:N, MBC, net NH_4_^+^ production rate, net nitrification rate, % cover of N-fixing species, % clay, bacterial abundance, and fungal abundance), on NO and N_2_O fluxes, we identified 11 models with delta < 2 for NO and 15 models with delta < 2 for N_2_O. When models with delta < 2 were averaged for NO and N_2_O, each averaged model contained eight final explanatory variables (**Figure 6; Table S4**). The large number of non-significant variables in the final models reflects high variation in the response (NO and N_2_O flux). NO was positively correlated with % clay (coefficient estimate 0.02 ± 0.008; p = 0.02) and negatively correlated with bacterial (coefficient estimate -4.59 E-10 ± 1.76 E-10; p = 0.01) and fungal abundance (coefficient estimate -2.97 E-09 ± 1.12 E-09; p = 0.01). N_2_O flux was positively correlated with GWC measured at the end of the soil incubations (coefficient estimate 1.5 ± 0.66; p = 0.03) and with % clay (coefficient estimate 0.04 ± 0.02; p = 0.02) but negatively correlated with MBC (coefficient estimate -0.002 ± 6.6 E-04; p = 0.007).

### 3.6 Natural Abundance Isotopes of N_2_O

The natural abundance N_2_O isotopocule values measured from high-emitting soils fell largely within or between literature-derived ranges for nitrification, bacterial denitrification, and fungal denitrification (**Table S5**) without one single process dominating as a source (**Figure 7**; **Table S6**). Site preference ranged between 4.2 and 47.5 ‰, consistent with values associated with contributions of denitrifying bacteria (low SP samples) and fungal denitrification and/or nitrifying bacteria (high SP samples; **Figure 7A**). Bulk δ^15^N_2_O ranged between -44 and -10.8 ‰, with three out of five samples falling along the expected N_2_O-to-N_2_ reduction line (Yu et al., 2020). Bulk δN_2_^18^O ranged between 18.8 and 55 ‰, pointing to partial contributions from many processes including nitrification in addition to bacterial and fungal denitrification (**Figure 7B**).

**Figure 7.**
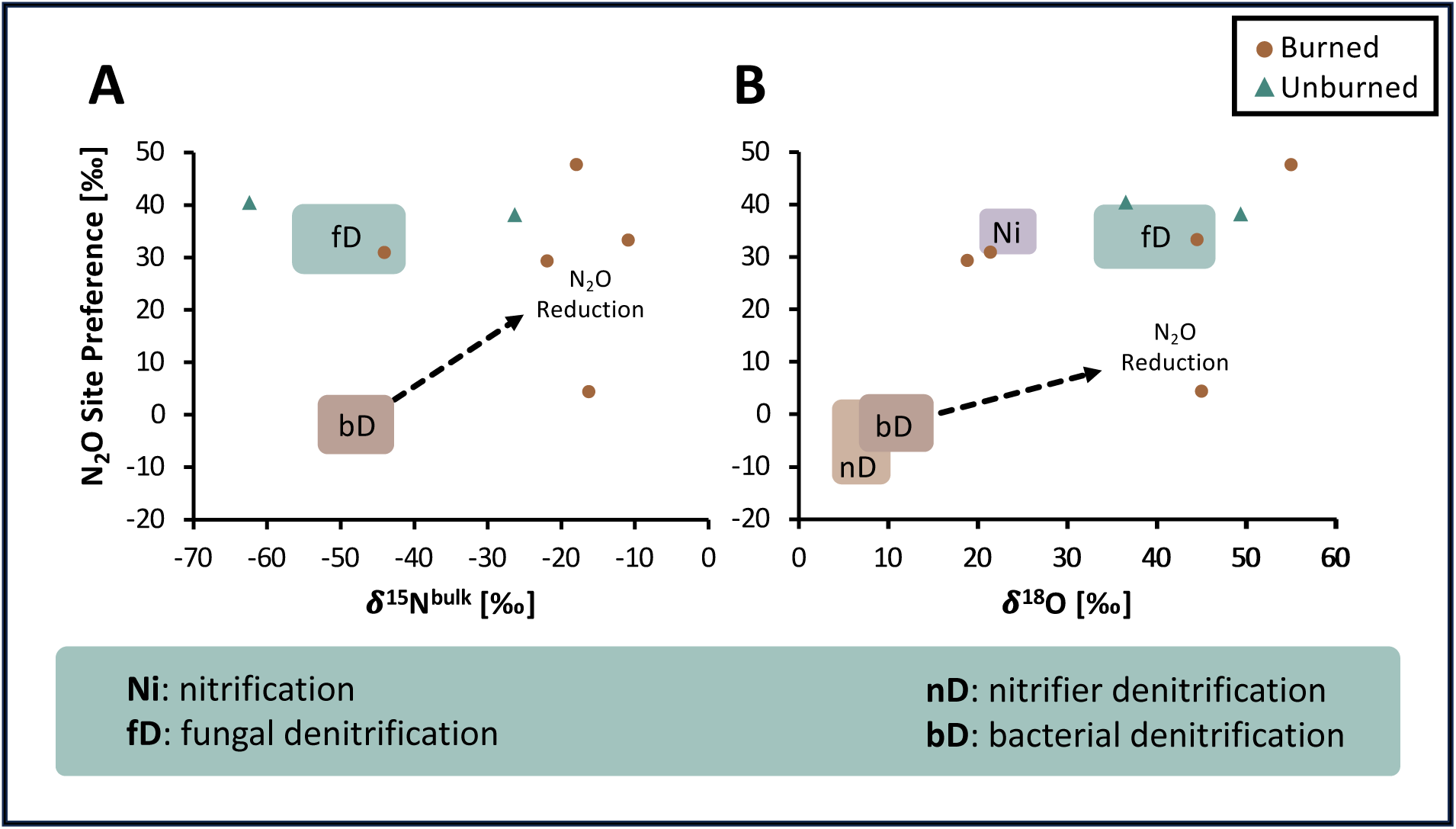
Natural abundance isotopic signature of N_2_O measured from six soil samples identified to have produced high concentrations of N_2_O over 48-h incubations. Boxes indicate literature-derived ranges for nitrification (Ni), nitrifier denitrification (nD), bacterial denitrification (bD), and fungal denitrification (fD) adjusted for average combined soil δ^15^N-NO_3_^-^ and δ^15^N-NO_2_^-^ (substrates assumed to contribute to denitrification processes bD and fD; **A**) and δ^18^O_H2O_ for nD, bD, and fD (**B**) according to recommendations by Yu et al. (2020; **Table S5**). The black arrow represents a line of expected isotope values associated with the reduction of N_2_O to N_2_.

When we estimated the fractional contributions of bacterial denitrification, nitrifier denitrification, fungal denitrification, and nitrifier nitrification using the FRAME model (Lewicki et al., 2022), we found a diversity of processes contributed to N_2_O fluxes post fire, including a reduction in the fraction of unreduced N_2_O, indicating increased N_2_O reduction to N_2_ (**Table S7**).

## 4. Discussion

We measured soil NO and N_2_O emissions along with a suite of biotic and abiotic variables to understand how wildfires alter these emissions across burn severities and over time. We found that wildfire increased soil NO emissions in post-fire years one and three, and while high variation in N_2_O emissions prevented detection of a burning effect, only burned soils produced N_2_O emission hotspots—N_2_O fluxes up to 165 × higher than the average in unburned plots and considered statistically high outliers. The highest fluxes of NO and N_2_O occurred in year one, corresponding to the highest soil extractable NH_4_^+^ and NO_3_^-^ availabilities, but the fluxes declined over time, supporting our hypothesis that burning would initially increase soil NO and N_2_O emissions but decrease as N substrates decreased (i). We also found that the highest fluxes of NO and N_2_O were measured in soils that burned at Medium severity, supporting our second hypothesis that Medium severity burning would create an optimal balance between high soil N availability and microbial survival to support high fluxes (ii). We also hypothesized that soil NO and N_2_O emissions would be positively correlated with pH, NH_4_^+^, NO_3_^-^, and microbial biomass (iii), but found that fluxes were positively related to abiotic variables such as % clay and soil GWC, while decreasing with increasing microbial biomass and abundance, suggesting that both soil chemistry and microbial recovery contribute to soil NO and N_2_O emissions. Overall, accelerated NO emissions and increased variability of N_2_O fluxes post fire indicate that wildfires can promote burn-severity-sensitive gaseous N losses long after they are extinguished due to interactions with post-fire soil chemistry and microbial biomass.

### 4.1 Soil NO and peak N_2_O emissions increased post fire, declining with soil N pools over time

Wildfire stimulated soil NO emissions and promoted N_2_O emission hotspots by initially increasing soil N availability but declined over time as substrates were exhausted. The highest NO emissions occurred in year one post fire (**Figure 2**), with peak fluxes more than three times the highest fluxes measured from a similar chaparral (unburned) site with comparably high atmospheric N deposition (Krichels et al., 2024). This strong increase in NO flux post fire is in line with previous work in chaparral, which found up to 300% increases in NO emissions in the first 6 months after wildfire (Levine et al., 1988).While soil N_2_O fluxes varied greatly in our study with no significant changes detectable post fire (**Figure 2C**), we observed several high outliers for N_2_O that occurred only in burned plots, with the highest measured in year one (∼165 × the average in unburned plots). Increases in NO and peak N_2_O emissions in burned plots corresponded to peak soil extractable NH_4_^+^ and NO_3_^-^ availability leading up to the end of year one (**Figure 3**), suggesting that the post-fire nutrient flush from ash deposition stimulated N emissions.

As both soil extractable NH_4_^+^ and NO_3_^-^ pools returned to unburned levels between years one and two, NO and peak N_2_O gas fluxes also decreased in year two. Similarly, bulk soil N decreased in year two, which when paired with the significant δ^15^N enrichment of soils from burned plots, suggests N cycling rates were high in the first years post fire, with high soil N losses leading to the observed isotopic enrichment of the residual N pool (**Figure S4B**; Deb et al., 2024; Liao et al., 2021). While we observed lower NO emissions as substrates decreased in year two, we observed a significant uptick in NO emissions from burned soil in the third post-fire year (**Figure 2A**). N-fixing plants have been linked to increased soil N availability and N emissions (Erickson & Perakis, 2014; Kou-Giesbrecht et al., 2021), and although only significant at p = 0.08, we saw an increase of N fixing plants over time in burned plots (**Figure S6**) that could have contributed to increased NO in year three. These findings suggest that high rates of post-fire N cycling and N gas emissions could be sustained over the first year after fire, dipping with the initial exhaustion of post-fire N substrates, but over longer time scales, N gas emissions may exhibit non-linear trends related to ecosystem recovery not previously captured for chaparral (previously measured only up to six months; Anderson et al., 1988; Anderson & Poth, 1989; Levine et al., 1988).

### 4.2 Wildfires stimulate nitrification and promote denitrification hotspots

Post-fire increases in soil NH_4_^+^ availability are suspected to stimulate nitrification and, thereby, contribute to high post-fire NO emissions (NO is typically the major by-product of nitrification; Firestone & Davidson, 1989). Previous evidence for high nitrification rates driving high emissions of NO in chaparral include NO:N_2_O ratios >1 typical of nitrification (Anderson et al., 1988; Anderson & Poth, 1989; Levine et al., 1988), a pattern we also observed despite lack of statistical significance (**Table 1**). Consistent with this, we found a 9-month lag between initially high soil NH_4_^+^ concentrations post fire and peak extractable NO_3_^-^, suggesting that nitrifying organisms oxidized NH_4_^+^ to NO_3_^-^ over time (**Figure 3**). This matches observed post-fire increases in the abundances of genes involved in nitrification pathways in the first year at our site (Pulido Barriga et al., 2025). This is also consistent with significantly more negative *in situ* net NH_4_^+^ production rates (i.e., the difference between gross NH_4_^+^ production and consumption rates) in burned soils (**Figure 4A**), suggesting that NH_4_^+^ was being consumed at greater rates after fire, in line with high nitrification activity. Furthermore, δN_2_^18^O and site preference values for two of the highest N_2_O emitting soils fell in the literature-derived ranges for nitrification in year one (**Figure 7B**), adding to existing support for the role of nitrification as an important driver of gaseous soil N emissions in post-fire environments.

High post-fire rates of nitrification, leading to a buildup of NO_3_^-^, are conditions well-known to favor denitrification (Anderson & Levine, 1986; Firestone & Davidson, 1989).

However, despite the high NO_3_^-^ availability leading up to the first-year sampling post fire (**Figure 3B**), we saw no statistical change in mean soil N_2_O emissions (**Figure 2C**). The high variability we observed in N_2_O emissions is consistent with variation common across soil microsites (Schlüter et al., 2024) and with genes encoding for denitrification pathways being rare relative to gene abundances for assimilatory and dissimilatory nitrate reductases within the first year post fire at our site (Pulido Barriga et al., 2025). Since denitrifiers need access to both N and C substrates (Knowles, 1982), the decrease in particulate and mineral-associated organic C observed at this site after the wildfire (Krichels et al., 2025) may have limited denitrification despite high N availability. However, the high N_2_O fluxes we observed only in burned soils suggests that wildfires may increase N availability in microsites to favor “hotspots” of denitrification, a phenomena previously observed in a mediterranean macchia shrubland (Fierro & Castaldi, 2011) and an Alaskan boreal forest (Morishita et al., 2015). This is consistent with our measurements of site preference and δ^15^N_2_O_bulk_ of high N_2_O-emitting soils matching literature-derived ranges for fungal and bacterial denitrification (**Figure 7A**; **Table S5**).

Moreover, we found that the GWC retained in soil microcosms at the end of the incubations and % clay positively correlated with N_2_O fluxes (**Figure 6B**), pointing to the importance of suboxic conditions for denitrification activity (Keiluweit et al., 2018). Clay can also provide chemically reactive surfaces and improve the availability of metal species that favor chemodenitrification (abiotic production pathway for NO and N_2_O; Heil et al., 2016). However, we suspect denitrification more strongly contributed to these post-fire hotspots as we found evidence for N_2_O reduction to N_2_, with several of the high N_2_O-emitting burned soils falling near the expected N_2_O to N_2_ reduction line (**Figure 7**) and four out of five soils showing decreases in residual N_2_O fractions pointing to increased reduction of N_2_O to N_2_ in FRAME estimates (N_2_ is the end product of denitrification; **Table S7**; Yu et al., 2020). Wildfire could promote the complete reduction of N_2_O to N_2_ by depositing wood char onto soils—a compound chemically similar to biochar known to facilitate complete denitrification and reduce N_2_O emissions by ∼40% (Cayuela et al., 2013; Edwards et al., 2018; Kaur et al., 2023). High N_2_ emissions have been observed in dry shrublands after fire (Dannenmann et al., 2018; Karhu et al., 2015), and high rates of N_2_O reduction are consistent with the abundance of low and negative individual N_2_O flux measurements in our dataset. Altogether, wildfire may exacerbate soil N_2_O flux variability by promoting complete reduction to N_2_ in some microsites and simultaneously generating hotspots in others.

### 4.3 Post-fire soil NO and N_2_O emissions were promoted at medium and high severities

Although our plots were characterized as Moderate burn severity at the 30 m^2^ scale by USFS BAER teams, we found that heterogeneity in soil burn severity at smaller scales can drive strong variation in N gas emissions. Wildfire promoted mean NO and N_2_O hotspot emissions at Medium and High soil burn severity categories (1-m^2^ scale), but NO emissions did not differ from unburned in the Low severity category (**Figure 2B**), and no hotspot emissions for N_2_O were measured at Low severity (**Figure 2D**). The increase in NO and N_2_O flux we observed at Medium and High severity in chaparral may help contextualize the strong fire response of shrublands (which tend to burn at Medium to High severities; Baeza et al., 2005; Barro & Conard, 1991) globally compared to other major ecosystem types: for example, grassland fires tend to burn at lower severities and produce lower NO and N_2_O fluxes post fire compared with shrublands and forests, whereas forests that can burn at a range of severities often produce highly variable fluxes (Stephens & Homyak, 2023).

We expected that soil N gas emissions would be highest at Medium soil burn severity due to the combination of higher pH and NH_4_^+^ relative to Low severity, and higher microbial survival relative to High severity. Increases in pH and NH_4_^+^ have been associated with increases in the abundance and activity of ammonia-oxidizing bacterial communities, which are linked to producing more NO and N_2_O relative to ammonia-oxidizing archaea (Ball et al., 2010; Mushinski et al., 2019; Prosser et al., 2019). We found that post-fire soil NH_4_^+^ and NO_3_^-^ concentrations were highest in the High category (**Table S3**; **Figure S3A&B**), and soil pH was significantly increased over unburned at both Medium and High burn severity categories (**Table S3**; **Figure S3C**), suggesting that the major changes to soil physicochemical properties occurred mainly at Medium and High severity to drive increases in NO and N_2_O emissions.

In our soils, microbial biomass C, bacterial, and fungal abundances were generally decreased by fire regardless of burn severity (**Figure S3D**, **Figure S5 B&D**), and when we examined drivers of NO and N_2_O fluxes across all severity categories (including unburned), we found that NO significantly decreased with increasing bacterial and fungal abundances and N_2_O decreased with increasing microbial biomass C (**Figure 6**). The decrease in N gas emissions with increasing microbial biomass and abundances suggests that other microbial processes such as N assimilation or dissimilatory nitrate reduction to ammonium (DNRA; a process whose genetic potential was found to increase after fire at this site and was consistent with observed production of NH_4_^+^ nine months post fire; Pulido Barriga et al., 2025) may compete with nitrifiers and denitrifiers for post-fire N resources. However, this trend did not directly relate to soil burn severity as expected and rather reflects the strong overall decrease in microbial biomass across all burn severity categories relative to unburned. Therefore, soil burn severity may more directly relate to changes in soil properties (pH, N pools) that influence NO and N_2_O emissions at landscape scales. These findings suggest that despite high variation at small scales, soil burn severity provides broad insight into changes in soil properties that influence mean NO and peak N_2_O emissions and may prove a valuable tool for large-scale assessment of the effects of wildfire on air quality and climate.

## 5. Conclusion

Wildfire increased soil emissions of NO for up to three years post fire, driven by changes in soil pH, soil NH_4_^+^ availability, and microbial biomass. Soil emissions of N_2_O did not differ significantly after fire due to high variation; however, only burned soils produced N_2_O fluxes that were up to ∼165 × higher than the average in unburned soils. Analysis of the isotopic composition of N_2_O from these high N_2_O-emitting soils suggests a diversity of contributing microbial processes, including nitrification and fungal and bacterial denitrification, pointing to increased heterogeneity in the post-fire environment that may generate hotspots of N_2_O production. Plot-level burn severity categories revealed elevated NO emissions and high N_2_O emission outliers occurred in Medium and High severity categories only, possibly indicating unique impacts from higher-severity wildfires relative to lower severity fires. Our study extends the timeframe over which post-fire soil N gas emissions have been measured in chaparral, provides evidence of non-linear trends in soil N emissions related to ecosystem recovery, and shows that wildfires increase soil emissions of trace N gases for years after the fire is extinguished by: i) increasing soil N availability, ii) shifting plant and microbial communities, and iii) increasing landscape heterogeneity to promote N_2_O hotspots which could scale up to affect regional air quality and global greenhouse gas balance.

## Acknowledgments

This research was supported by the California Department of Forestry and Fire Protection (award 8GG20812 to E.Z.S), the US National Science Foundation (DEB 1916622 to P.M.H), US Department of Agriculture NIFA award (2022-67014-36675 to S.I.G and P.M.H), US Department of Energy BER award (DE-SC0023127 to S.I.G and P.M.H), and US NSF Graduate Research Fellowship Program (award #DGE-1326120 to MK). This work was supported in part by the USDA Forest Service Rocky Mountain Research Station. The findings and conclusions in this publication are those of the authors and should not be construed to represent any official USDA or U.S. Government determination or policy. We thank the Cleveland National Forest and the Trabuco Ranger Districts, especially Darrel Vance, Emily Fudge, Jeffrey Heys, Lauren Quon, Jacob Rodriguez, and Victoria Stempniewicz for their guidance and help with permitting and site selection. We thank James Randolph, Dylan Enright, Sameer S. Saroa, Taylor Naquin, Judy A. Chung, Aishwarya Veerabahu, and Arik Joukhajian, for help in the field and with vegetation and microbial work, and we thank Sharon Zhao, Johann Püspök, Melissa Zavala, Yareli Olazabal, Tony Calma, Nikki Shelton, Heather Haro, Kobe Luu, Karen Argumedo, and David Lyons at the Environmental Sciences Research Laboratory for their assistance with biogeochemical analyses.

