## Supplementary Materials for "Wildfire alters nitrogen cycling to increase soil emissions of nitric oxide (NO) and the heterogeneity of nitrous oxide (N_2_O) in California chaparral"

**Supplementary Tables**

| Plot | Treatment | Latitude | Longitude | Elevation | Burn Severity (BAER) | Average Ash depth (cm) | Bulk Density | % Clay | % Sand | % Silt | Soil Series and Taxonomic Classification |
|---|---|---|---|---|---|---|---|---|---|---|---|
| 1 | Burned | 33.6901 | -117.463 | 1288 | Moderate | 4.8 | 0.90 | 3.8 | 76.1 | 20.1 | Cieneba series; Loamy, |
| 2 | Burned | 33.69537 | -117.471 | 1260 | Moderate | 5.5 | 0.86 | 8.4 | 74.3 | 17.4 | mixed, superactive, |
| 3 | Burned | 33.69326 | -117.467 | 1237 | Moderate | 2.9 | 0.79 | 13.1 | 54.3 | 32.6 | nonacid, thermic, shallow |
| | | | | | | | | | | | Typic Xerorthents |
| 4 | Burned | 33.68456 | -117.457 | 1260 | Moderate | 2.6 | 0.95 | 15.4 | 30.2 | 54.4 | Friant series; Loamy, |
| 5 | Burned | 33.67809 | -117.457 | 1195 | Moderate | 6.5 | 0.82 | 11.6 | 40.3 | 48.1 | mixed, superactive, |
| 6 | Burned | 33.67168 | -117.459 | 1285 | Moderate | 5.2 | 0.59 | 11.1 | 45.9 | 43.0 | thermic Lithic |
| 7 | Unburned | 33.67135 | -117.459 | 1283 | | | 0.49 | 8.7 | 45.9 | 45.4 | Haploxerolls |
| 8 | Unburned | 33.66813 | -117.456 | 1250 | | | 0.61 | 14.8 | 48.2 | 37.0 | |
| 9 | Unburned | 33.6678 | -117.455 | 1240 | | | 0.58 | 19.2 | 37.7 | 43.1 | |

**Table S1.** Soil physical properties and taxonomic classification at sampling plots. Soil bulk density (g $cm^{-3}$), % sand, % clay, and % silt were averaged over 4 subplots in each plot. Burn severity is reported as plot-level classifications derived from the Holy Fire Burned Area Emergency Report assessment by the US Forest Service at a resolution of 30 $m^2$. Soil series were mapped at the finest available scale with the USDA National Cooperative Survey Web Soil Survey online tool.

| **Table S2** | | **Burned** | | | **Unburned** | | | **Estimate** | **P-values** | | |
|---|---|---|---|---|---|---|---|---|---|---|---|
| **Variable (*in situ*)** | unit | Mean | SE | n | Mean | SE | n | Burn | Burn | Time | Burn x Time |
| Bulk Density | g $cm^{-3}$ | 0.79 | 0.05 | 12 | 0.56 | 0.05 | 12 | 0.23 | **0.004** | NA | NA |
| Soil Moisture | g water g $soil^{-1}$ | 0.026 | 0.002 | 36 | 0.052 | 0.006 | 36 | -0.026 | **0.025** | 0.334 | 0.122 |
| pH | | 6.54 | 0.06 | 36 | 6.13 | 0.05 | 35 | 0.39 | **<0.001** | 0.471 | 0.103 |
| Bulk soil C | mg C g $soil^{-1}$ | 27.34 | 1.63 | 72 | 46.26 | 7.26 | 18 | -22.06 | **0.007** | 0.225 | **0.001** |
| Bulk soil N | mg N g $soil^{-1}$ | 1.45 | 0.09 | 72 | 1.87 | 0.26 | 18 | -0.44 | 0.183 | **0.046** | **0.003** |
| C:N | ratio | 22.6 | 0.8 | 36 | 27.8 | 1.2 | 18 | -4.7 | **0.028** | **<0.001** | 0.627 |
| Bulk soil $\delta^{15}N$ | ‰ | 2.42 | 0.15 | 36 | 1.80 | 0.38 | 18 | 0.60 | 0.118 | 0.087 | **0.009** |
| Bulk soil $\delta^{13}C$ | ‰ | -26.39 | 0.16 | 36 | -26.87 | 0.35 | 18 | 0.50 | 0.176 | **<0.001** | **0.030** |
| Ammonium | µg $NH_4^+$-N g $soil^{-1}$ | 21.2 | 2.7 | 36 | 5.8 | 2.5 | 36 | 15.0 | **0.001** | **0.002** | **0.002** |
| Nitrate | µg $NO_3^-$-N g $soil^{-1}$ | 13.8 | 2.8 | 36 | 0.7 | 0.2 | 36 | 13.3 | **<0.001** | **<0.001** | **<0.001** |
| MBC | µg C g $soil^{-1}$ | 145.7 | 13.0 | 36 | 370.8 | 50.3 | 36 | -220.5 | **0.002** | **0.001** | **0.043** |
| MBN | µg N g $soil^{-1}$ | 6.1 | 1.2 | 36 | 21.9 | 3.5 | 36 | -15.8 | **0.002** | 0.247 | 0.290 |
| Bacterial abundance | gene copy number | 2.06E+08 | 1.34E+07 | 36 | 5.57E+08 | 2.69E+08 | 35 | -3.20E+08 | 0.248 | 0.162 | 0.248 |
| Fungal abundance | gene copy number | 1.55E+07 | 1.65E+06 | 36 | 7.80E+07 | 8.61E+06 | 35 | -6.21E+07 | **<0.001** | **0.024** | 0.478 |
| Bacterial Richness | Total no. bacterial sp. | 555 | 48 | 35 | 575 | 36 | 31 | -21 | 0.627 | **<0.001** | 0.253 |
| Fungal Richness | Total no. fungal sp. | 93 | 5 | 36 | 286 | 15 | 35 | -194 | **<0.001** | **0.042** | 0.484 |
| N-fixing plants | % cover N fixing sp. | 6.0 | 2.6 | 36 | 3.2 | 1.0 | 36 | 3.1 | 0.346 | **0.016** | **0.059** |
| Vegetation Richness | Total no. plant species | 4 | 0.4 | 36 | 2 | 0.2 | 18 | 1.4 | 0.105 | **0.008** | 0.221 |
| **Variable (Laboratory)** | | Mean | SE | n | Mean | SE | n | Burn | Burn | Time | Burn x Time |
| $CO_2$ flux | ng $CO_2$-C $g^{-1}$soil | 194301 | 17463 | 36 | 170621 | 23694 | 18 | 23813 | 0.227 | 0.606 | 0.303 |
| NO flux | ng NO-N $g^{-1}$soil | 177.9 | 26.9 | 36 | 87.4 | 15.4 | 18 | 90.6 | **0.007** | 0.536 | **0.049** |
| $N_2O$ flux | ng $N_2O$-N $g^{-1}$soil | 295.7 | 163.7 | 36 | 30.9 | 17.4 | 18 | 265.7 | 0.095 | 0.345 | 0.100 |
| NO:$N_2O$ | ratio | 17.0 | 19.1 | 72 | -0.1 | 2.1 | 18 | 4.1 | 0.922 | 0.217 | 0.937 |
| Net mineralization | µg N g $soil^{-1}$ $day^{-1}$ | 7.06 | 2.84 | 36 | 3.55 | 1.03 | 17 | 3.55 | 0.348 | **0.002** | **0.051** |
| Net nitrification | µg $NO_3^-$-N g $soil^{-1}$ $day^{-1}$ | 6.94 | 2.86 | 36 | 0.25 | 0.19 | 17 | 6.67 | 0.087 | **0.002** | **0.041** |
| Net $NH_4^+$ production | µg $NH_4^+$-N g $soil^{-1}$ $day^{-1}$ | 0.12 | 0.98 | 36 | 3.30 | 1.02 | 17 | -2.62 | 0.193 | 0.062 | 0.644 |

**Table S2**. Summary of differences between mean burned and mean unburned using only Mollisols measured at the end of years one, two, and three post fire (directly comparable to **Table 1** which includes both Mollisols and Entisols**)**. “Estimate” is the average unit change in burned compared to unburned plots over all three years combined, while p-values show interactions with time modeled with linear mixed effects models. Bulk density was measured at only one sampling time, so no time interactions are shown.

| Table S3 | Unburned | | | Low | | | Medium | | | High | | |
|---|---|---|---|---|---|---|---|---|---|---|---|---|
| **Variable (*in situ*)** | Mean | SE | n | Mean | SE | n | Mean | SE | n | Mean | SE | n |
| Bulk Density | 0.56 | 0.05 | 12 | 0.84 | 0.06 | 3 | 0.72 | 0.11 | 4 | 0.84 | 0.04 | 17 |
| Soil Moisture | 0.052 | 0.006 | 36 | 0.016 | 0.002 | 11 | 0.022 | 0.003 | 12 | 0.021 | 0.002 | 49 |
| pH | 6.13 | 0.05 | 35 | 6.43 | 0.21 | 11 | 6.69 | 0.09 | 12 | 6.58 | 0.06 | 49 |
| Bulk soil C | 46.26 | 7.26 | 18 | 18.65 | 2.63 | 11 | 31.06 | 2.75 | 12 | 28.38 | 2.14 | 49 |
| Bulk soil N | 1.87 | 0.26 | 18 | 1.05 | 0.19 | 11 | 1.64 | 0.14 | 12 | 1.49 | 0.12 | 49 |
| C:N | 27.8 | 1.2 | 18 | 21.4 | 1.0 | 11 | 22.3 | 1.0 | 12 | 22.6 | 0.6 | 49 |
| Bulk soil $\delta^{15}N$ | 1.80 | 0.38 | 18 | 2.28 | 0.37 | 11 | 2.44 | 0.32 | 12 | 1.98 | 0.16 | 49 |
| Bulk soil $\delta^{13}C$ | -26.87 | 0.35 | 18 | -26.43 | 0.18 | 11 | -27.09 | 0.24 | 12 | -26.56 | 0.12 | 49 |
| Ammonium | 5.8 | 2.5 | 36 | 12.5 | 2.1 | 11 | 13.4 | 3.1 | 12 | 17.6 | 2.4 | 49 |
| Nitrate | 0.7 | 0.2 | 36 | 4.5 | 2.2 | 11 | 9.8 | 4.1 | 12 | 9.9 | 2.2 | 49 |
| MBC | 370.8 | 50.3 | 36 | 119.9 | 28.2 | 11 | 156.1 | 14.7 | 12 | 133.4 | 12.6 | 49 |
| MBN | 21.9 | 3.5 | 36 | 4.8 | 1.6 | 11 | 5.8 | 2.0 | 12 | 5.0 | 0.9 | 49 |
| Bacterial abundance | 5.57E+08 | 2.69E+08 | 35 | 1.26E+08 | 2.68E+07 | 9 | 2.36E+08 | 2.35E+07 | 12 | 1.68E+08 | 1.15E+07 | 49 |
| Fungal abundance | 7.80E+07 | 8.61E+06 | 35 | 9.91E+06 | 2.18E+06 | 9 | 1.62E+07 | 2.26E+06 | 12 | 1.62E+07 | 2.48E+06 | 49 |
| Bacterial Richness | 575 | 36 | 31 | 598 | 113 | 10 | 572 | 73 | 11 | 531 | 35 | 49 |
| Fungal Richness | 286 | 15 | 35 | 129 | 19 | 9 | 120 | 9 | 12 | 94 | 6 | 49 |
| N-fixing plants | 3.2 | 1.0 | 36 | 9.2 | 4.7 | 11 | 1.8 | 0.9 | 12 | 5.9 | 1.9 | 49 |
| Vegetation Richness | 2 | 0.2 | 18 | 4 | 0.9 | 11 | 3 | 0.6 | 12 | 4 | 0.3 | 49 |
| **Variable (Laboratory)** | Mean | SE | n | Mean | SE | n | Mean | SE | n | Mean | SE | n |
| $CO_2$ flux | 170621 | 23694 | 18 | 136936 | 27689 | 11 | 166678 | 17577 | 12 | 154446 | 14668 | 49 |
| NO flux | 87.4 | 15.4 | 18 | 192.8 | 90.9 | 11 | 208.4 | 70.2 | 12 | 142.9 | 12.3 | 49 |
| $N_2O$ flux | 30.9 | 17.4 | 18 | 459.2 | 453.1 | 11 | 430.7 | 404.4 | 12 | 121.5 | 72.8 | 49 |
| $NO:N_2O$ | -0.1 | 2.1 | 18 | -3.1 | 2.2 | 11 | 104.0 | 111.8 | 12 | 0.2 | 6.5 | 49 |
| Net mineralization | 3.55 | 1.03 | 17 | 0.67 | 2.80 | 9 | 8.76 | 6.25 | 12 | 4.41 | 2.16 | 49 |
| Net nitrification | 0.25 | 0.19 | 17 | 1.82 | 2.38 | 9 | 7.75 | 6.36 | 12 | 3.60 | 1.91 | 49 |
| Net $NH_4^+$ production | 3.30 | 1.02 | 17 | -1.15 | 1.04 | 9 | 1.02 | 4.57 | 12 | 0.81 | 0.67 | 49 |

**Table S3**. Means, standard error, and sample sizes for soil variables averaged across years one, two, and three at each burn severity category (unburned, low, medium, high) assessed at the $1m^2$ scale.

| **NO Burned Model Averages** | | | | | |
|---|---|---|---|---|---|
| **Variable** | **units** | **Estimate** | **SE** | **z value** | **p-value** |
| (Intercept) | | 1.77E+00 | 4.03E-01 | 4.358 | **<0.001** |
| GWC | g water g soil$^{-1}$ (end incubation) | 2.35E-01 | 3.17E-01 | 0.733 | 0.463 |
| % Clay | | 2.20E-02 | 8.91E-03 | 2.372 | **0.018** |
| pH | | 2.41E-02 | 5.74E-02 | 0.417 | 0.677 |
| MBC | Microbial biomass C: μg C g soil$^{-1}$ | 9.84E-09 | 2.28E-06 | 0.004 | 0.997 |
| $NH_4^+$ Production | μg N-$NH_4^+$ g soil$^{-1}$ day$^{-1}$ | 6.08E-06 | 2.24E-04 | 0.027 | 0.979 |
| N fixers | % cover of N fixing sp. | 2.62E-05 | 6.41E-04 | 0.040 | 0.968 |
| Bacteria | Abundance: gene copy number | -4.59E-10 | 1.76E-10 | 2.539 | **0.011** |
| Fungi | Abundance: gene copy number | -2.97E-09 | 1.12E-09 | 2.588 | **0.010** |
| **$N_2O$ Burned Model Averages** | | | | | |
| **Variable** | **units** | **Estimate** | **SE** | **z value** | **p-value** |
| (Intercept) | | 7.81E-01 | 6.97E-01 | 1.110 | 0.267 |
| GWC | g water g soil$^{-1}$ (end incubation) | 1.50E+00 | 6.56E-01 | 2.226 | **0.026** |
| % Clay | | 3.78E-02 | 1.57E-02 | 2.300 | **0.021** |
| pH | | 4.14E-02 | 1.00E-01 | 0.410 | 0.682 |
| C:N | | 2.66E-04 | 2.79E-03 | 0.094 | 0.925 |
| MBC | Microbial biomass C: μg C g soil$^{-1}$ | -1.82E-03 | 6.62E-04 | 2.686 | **0.007** |
| Nitrification | μg N-$NO_3^-$ g soil$^{-1}$ day$^{-1}$ | -3.84E-05 | 6.12E-04 | 0.062 | 0.950 |
| N fixers | % cover of N fixing sp. | -2.04E-04 | 2.48E-03 | 0.081 | 0.935 |
| Fungi | Abundance: gene copy number | -1.06E-17 | 2.23E-13 | 0.000 | >0.99 |

**Table S4.** Averaged model outputs of top models (delta <2) for soil NO and $N_2O$ emissions as shown in **Figure 6**. NO and $N_2O$ were log-transformed and modeled separately using a linear mixed effects multiple regression approach that included a random effect for plot, a temporal autocorrelation term, and controlled for variance between soil types. P-values show significance of averaged modeled estimates.

| Yu et al. (2020) | **$\delta^{15}N_2O_{bulk}$** | | **$\delta^{15}N_2O_{SP}$** | | **$\delta N_2{}^{18}O$** | |
|---|---|---|---|---|---|---|
| Process | low | high | low | high | low | high |
| bD | -52.8 | -42.4 | -7.5 | 3.7 | 7.7 | 14.3 |
| nD | | | -13.6 | 1.9 | 3.4 | 10.3 |
| Ni | | | 32 | 38.7 | 20.5 | 26.5 |
| fD | -54.5 | -39.5 | 27.2 | 39.9 | 33.0 | 46.1 |
| Average ‰ of soil substrates: | | $\delta^{15}N\text{-}NO_3^-$ | | $\delta^{18}O_{H2O}$ | | |
| | | -8.5 | | -9 | | |

**Table S5**. Literature-derived ranges for isotopic composition of $N_2O$ from nitrification (Ni), nitrifier denitrification (nD), bacterial denitrification (bD), and fungal denitrification (fD) derived from Yu et al. (2020) and Lewicki et al. (2022). Because these ranges assume a substrate contribution of 0 ‰, average combined soil $\delta^{15}N\text{-}NO_3^-$ and $\delta^{15}N\text{-}NO_2^-$ (substrates assumed to contribute to denitrification processes bD and fD) and $\delta^{18}O_{H2O}$ (assumed to contribute to nD, bD, and fD) were used to adjust these ranges according to recommendations by Yu et al. (2020) and Lewicki et al. (2022). We did not measure $\delta^{15}N\text{-}NH_4^+$ from these soils; therefore, we do not present adjusted ranges for Ni or nD which assume $NH_4^+$ to be the main substrate.

| | | | $N_2O$ (ppm) | | $NN^{15}O$ | | $N^{15}NO$ | | $NNO^{18}$ | |
|---|---|---|---|---|---|---|---|---|---|---|
| Plot ID | Burn | Year | Mean | SD | Mean | SD | Mean | SD | Mean | SD |
| 3S | Burned | 1 | 2.337 | 0.01 | 0.00853 | 6.75E-05 | 0.00834 | 7.17E-05 | 0.00489 | 4.75E-05 |
| 6W | Burned | 1 | 2.863 | 0.03 | 0.01035 | 1.10E-04 | 0.01013 | 1.08E-04 | 0.00591 | 6.94E-05 |
| 5W | Burned | 2 | 1.691 | 0.007 | 0.00613 | 4.16E-05 | 0.00609 | 4.79E-05 | 0.00352 | 4.38E-05 |
| 2N | Burned | 3 | 2.421 | 1.08 | 0.00884 | 3.93E-03 | 0.00856 | 3.76E-03 | 0.00510 | 2.25E-03 |
| 5S | Burned | 3 | 2.832 | 0.02 | 0.01010 | 7.62E-05 | 0.00987 | 8.07E-05 | 0.00585 | 5.59E-05 |
| 7E | Unburned | 1 | 2.173 | 0.01 | 0.00788 | 5.48E-05 | 0.00768 | 5.77E-05 | 0.00455 | 4.57E-05 |
| 7W | Unburned | 3 | 3.492 | 0.03 | 0.01229 | 1.04E-04 | 0.01191 | 1.03E-04 | 0.00729 | 7.25E-05 |

**Table S6**. Isotopocule concentrations for selected soils that produced the highest fluxes of $N_2O$ averaged over 3 minutes (n = 180) with associated standard deviations as a measure of uncertainty. Unburned soil from year 2 did not produce enough $N_2O$ (>1.2 ppm) to allow for isotopic analysis using our methods. Isotopocule concentrations were converted to delta values using equations 1-3 as displayed in **Figure 7**.

| Plot ID | Burn | Year | bD | nD | fD | Ni | r |
|---|---|---|---|---|---|---|---|
| 3S | burned | 1 | 0.31 ± 0.21 | 0.27 ± 0.19 | 0.22 ± 0.16 | 0.20 ± 0.15 | 0.03 ± 0.02 |
| 6W | burned | 1 | 0.32 ± 0.20 | 0.32 ± 0.20 | 0.16 ± 0.13 | 0.20 ± 0.14 | 0.05 ± 0.05 |
| 5W | burned | 2 | 0.41 ± 0.26 | 0.46 ± 0.26 | 0.07 ± 0.06 | 0.06 ± 0.05 | 0.06 ± 0.04 |
| 2N | burned | 3 | 0.18 ± 0.15 | 0.16 ± 0.14 | 0.33 ± 0.22 | 0.33 ± 0.21 | 0.04 ± 0.06 |
| 5S | burned | 3 | 0.14 ± 0.11 | 0.16 ± 0.11 | 0.13 ± 0.11 | 0.57 ± 0.18 | 0.50 ± 0.23 |
| 7E | unburned | 1 | 0.20 ± 0.15 | 0.18 ± 0.14 | 0.31 ± 0.22 | 0.32 ± 0.21 | 0.09 ± 0.05 |
| 7W | unburned | 3 | 0.14 ± 0.14 | 0.23 ± 0.18 | 0.12 ± 0.13 | 0.52 ± 0.26 | 0.10 ± 0.10 |

**Table S7**. Modeled fractional contributions of bacterial denitrification (bD), nitrifier denitrification (nD), fungal denitrification (fD), nitrifier nitrification (Ni), and the fraction of unreduced $N_2O$ (r) to soil $N_2O$ emissions (± standard deviation) from selected high-emitting soils estimated using the FRAME isotopic fractionation and mixing evaluation tool (https://malewick.github.io/frame/). Sources of $N_2O$ for each process are based on literature-derived values (Yu et al., 2020) and fractionation processes contributing to $N_2O$ reduction are based on experimental data from Well & Flessa (2010). Model structure and auxiliary inputs matched example 5.4 in Lewicki et al. (2022).

### Supplementary Figures

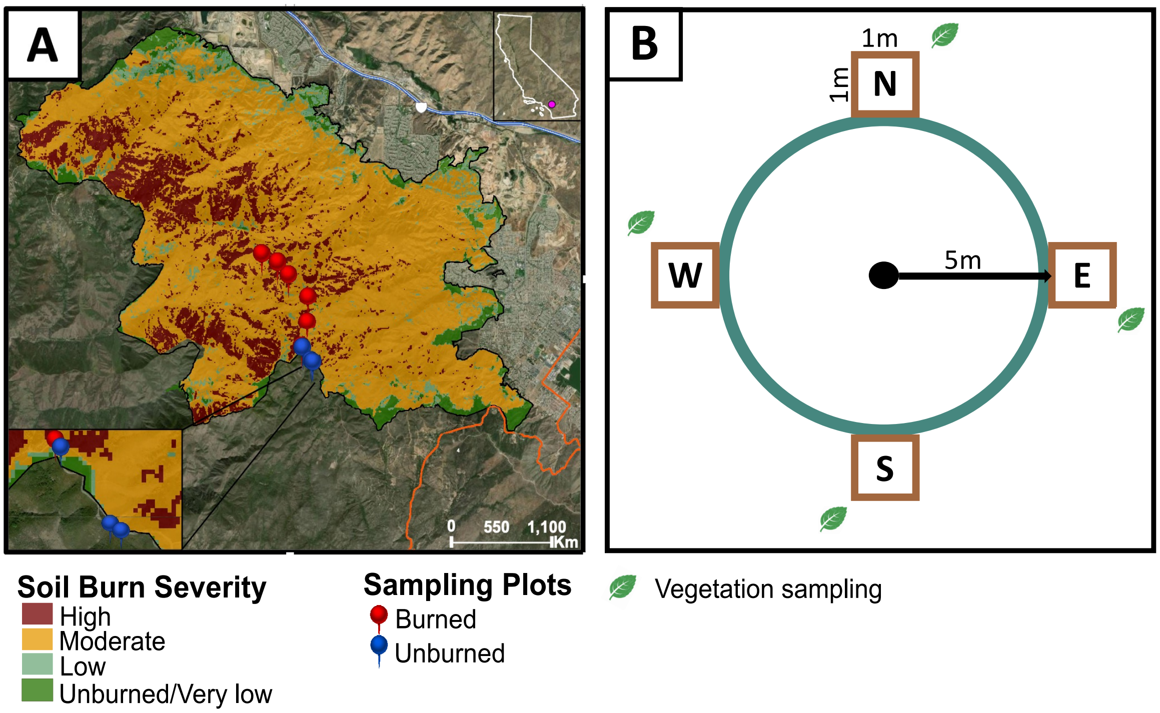


**Figure S1.** Panel **A** shows the extent of the Holy Fire, which burned 23,000 acres in the Cleveland National Forest, CA, USA, in September 2018 (black outline). Soil burn severity classifications are based on BAER (Burned Area Emergency Response) assessments performed by the US Forest Service. Pins show the location of 9 plots (red = burned, n = 6; blue = unburned, n = 3), which are each organized around a central plot with four 1 $m^2$ subplots 5 m from the center in each cardinal direction (**B**). Soil cores were taken down to 10 cm depth within each subplot, and vegetation was monitored in adjacent plots.

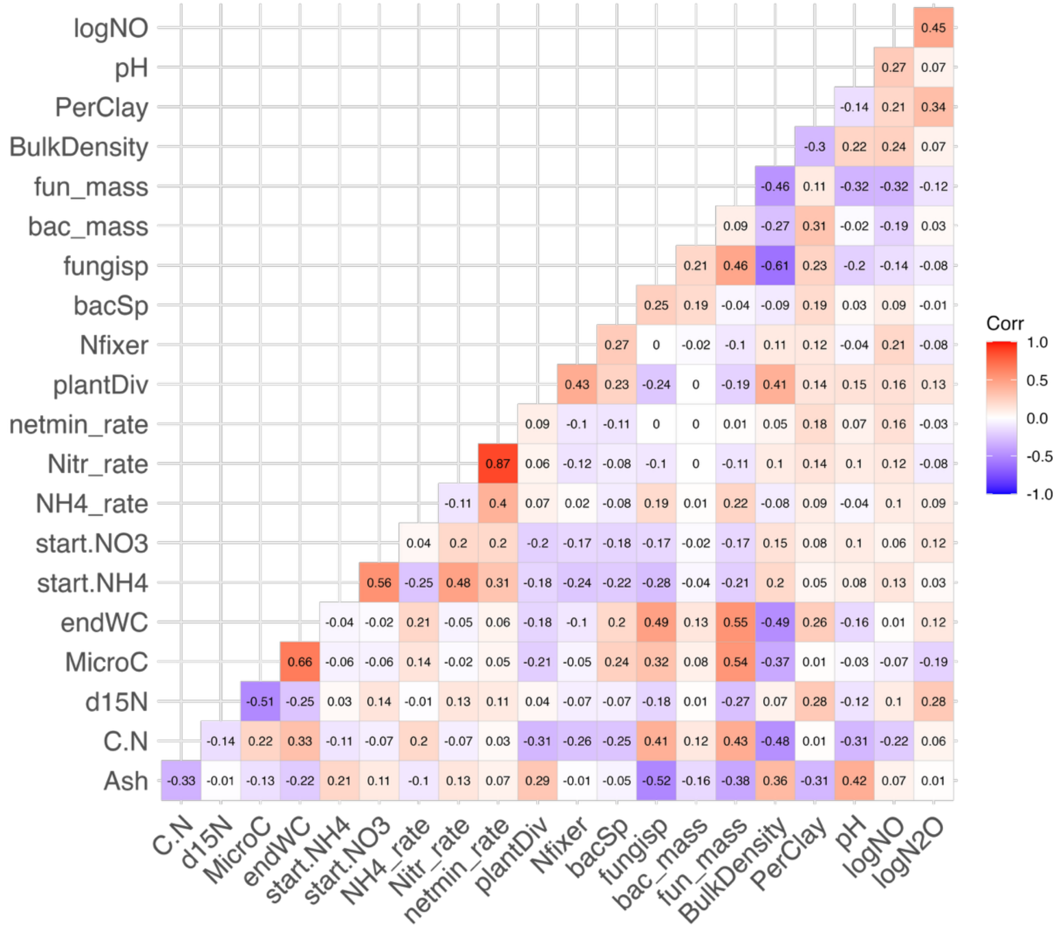


**Figure S2.** Pearson's correlations for all variables considered for candidates to include in the final global models for NO and $N_2O$. When variables had correlations > ± 0.7, only one was included to avoid collinearity in the models. Variable abbreviations represent the following: logNO and log$N_2O$ = the log-transformation of cumulative NO and $N_2O$ over 40 h; PerClay = % clay (chosen to represent texture due to co-linearity with % silt and % sand); BulkDensity = soil bulk density (g $cm^{-3}$); fun_mass and bac_mass = abundances of fungi (fun) and bacteria (bac); fungisp and bacsp = fungal and bacterial species richness; Nfixer = % cover of N-fixing plant species; plantDiv = species richness of plant community; netmin_rate = net N mineralization rate during gas flux incubation; Nitr_rate = net nitrification rate during gas flux incubation; NH4_rate

= net NH4 production rate during gas flux incubation; start.NO3 and start.NH4 = concentration (µg $NH_4^+$ /$NO_3^-$ -N g $soil^{-1}$) at beginning of gas flux incubation; endWC = gravimetric water content at the end of the gas flux incubation (all soils were wet to 75% water holding capacity at start, so this was chosen to represent different levels of drying or moisture retention during incubation); MicroC = microbial biomass C (µg C g $soil^{-1}$); d15N = bulk soil $\delta^{15}N$ (not chosen to include in models because this is not a driver of fluxes, but more likely to reflect high N cycling rates); C.N = bulk soil C:N ratio.

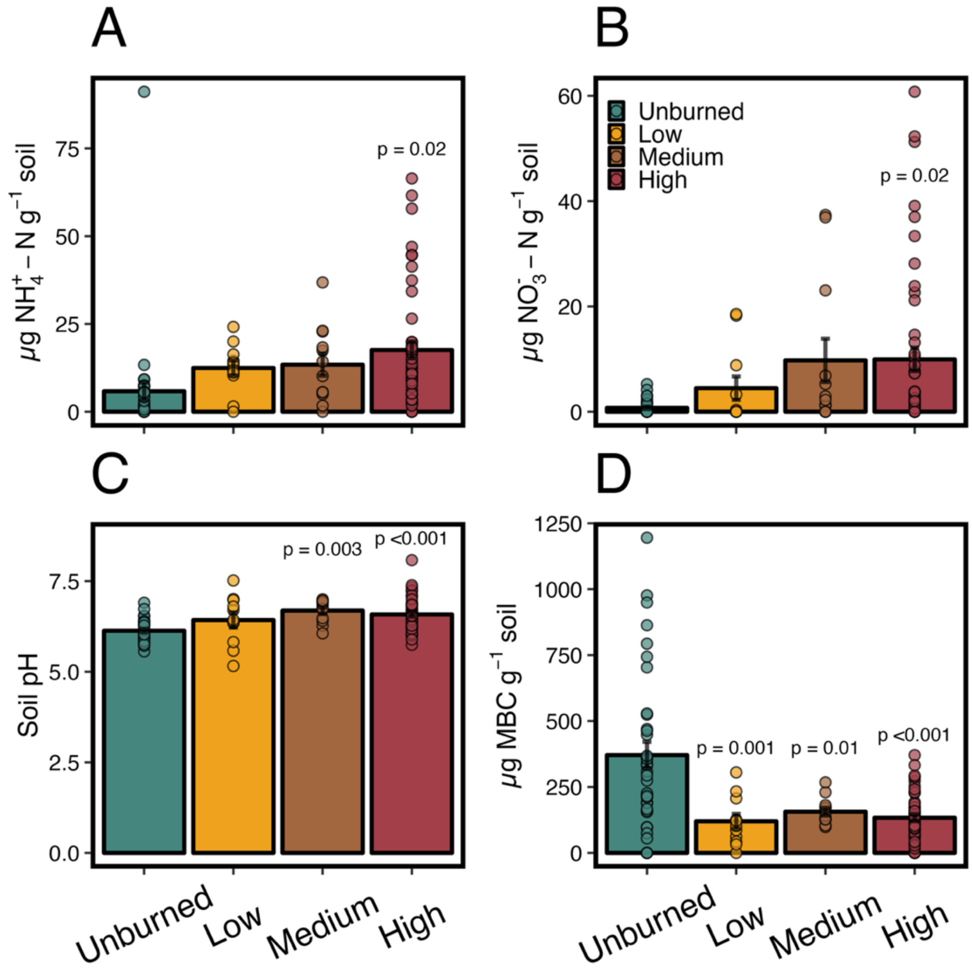


**Figure S3.** Variables measured at the ends of years 1–3 across fine-scale (1 $m^2$) soil burn severity categories: *in situ* soil extractable ammonium ($NH_4^+$; **A**), extractable nitrate ($NO_3^-$; **B**), soil pH (**C**), and microbial biomass C (MBC; **D**). Points represent means for each category, error bars are standard error, and samples sizes for each category can be found in **Table S3**. Linear mixed effects models were used estimate differences between burn severity categories, and p-values represent significant differences between individual burn severity categories compared to unburned. Post-hoc comparisons showed no differences between burned soils in Low, Medium, and High categories for **A**, **B**, **C**, or **D**.

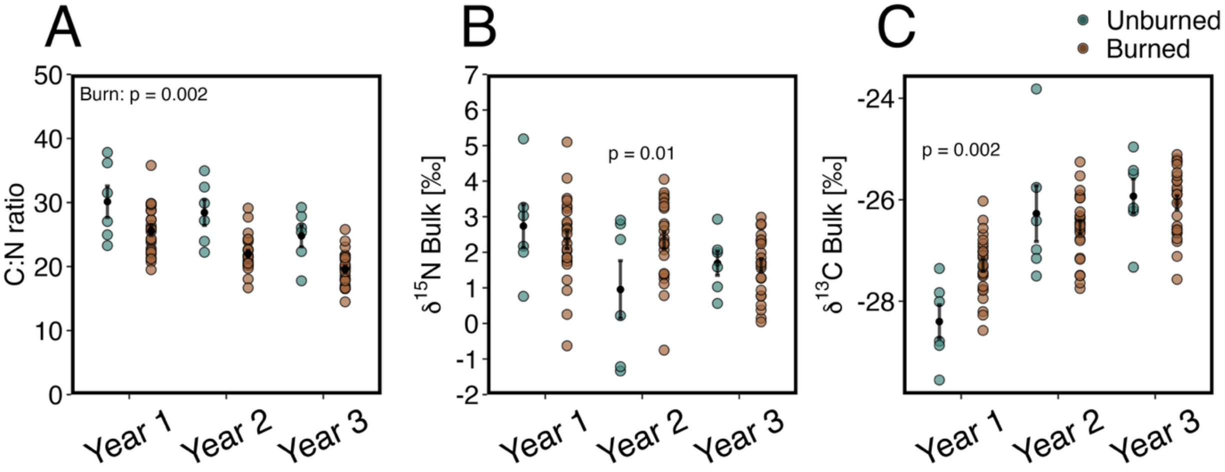


**Figure S4**. Mean bulk soil C:N ratios (**A**), bulk soil $\delta^{15}N$ (**B**), and bulk soil $\delta^{13}C$ (**C**) at each year of the study. Black dots represent means and error bars are standard error, for each timepoint n = 6 unburned, n = 24 burned. C:N ratios were consistently lower after fire (**A**; Burn: $p = 0.002$), but fire effects on bulk soil $\delta^{15}N$ and $\delta^{13}C$ differed only in individual years (p-values represent post-hoc analysis **B** and **C**).

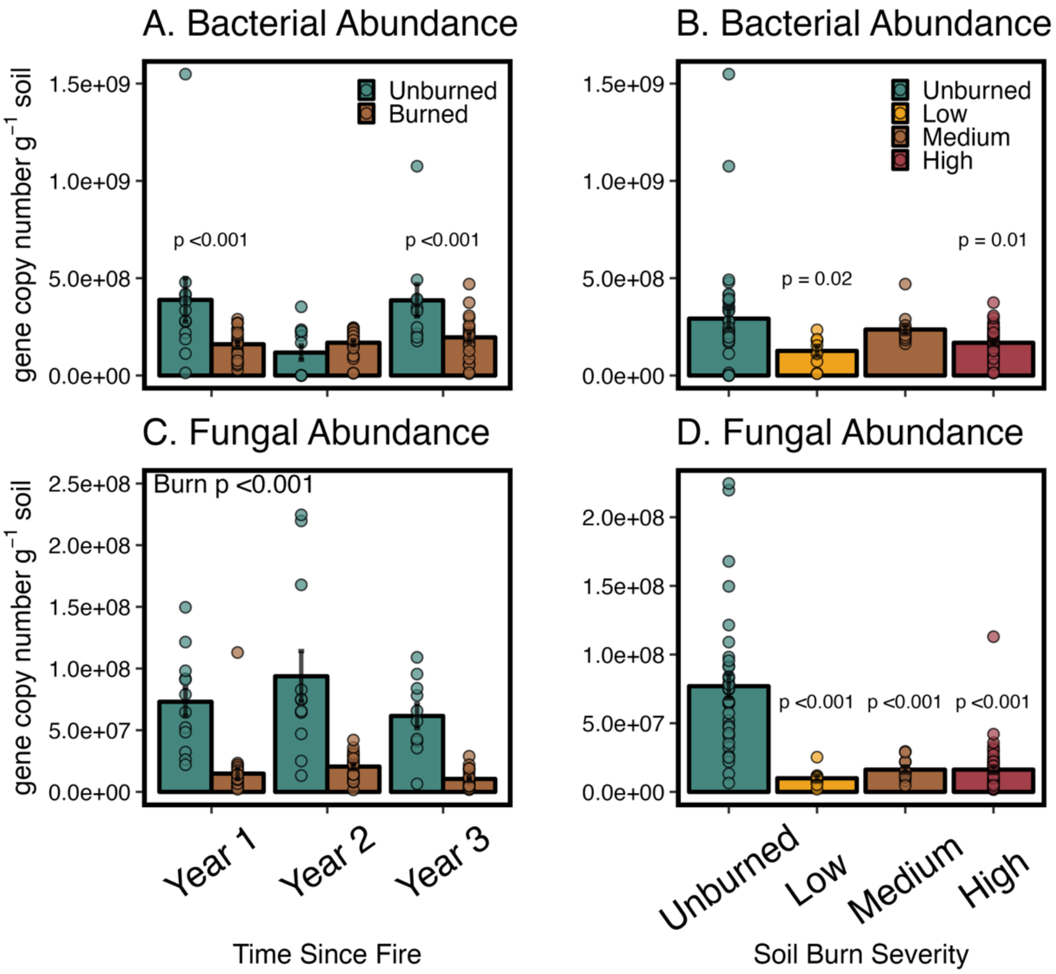


**Figure S5**. Mean gene copy numbers $g^{-1}$ soil for bacteria and fungi compared between burned and unburned (estimate of abundance; **A&C**) and between soil burn severity categories (**B&D**) at the end of years 1–3 post-fire. Error bars are standard error, for each timepoint n = 12 unburned, n = 24 burned. Dots represent individual observations. Where significant burn × time interactions were found, p-values are post-hoc comparisons of burned and unburned in each individual year (**A**), and "Burn" represents the overall effect of burning where no time interactions were present (**C**). P-values in **B** and **D** show significant differences between each burn severity category compared to unburned.

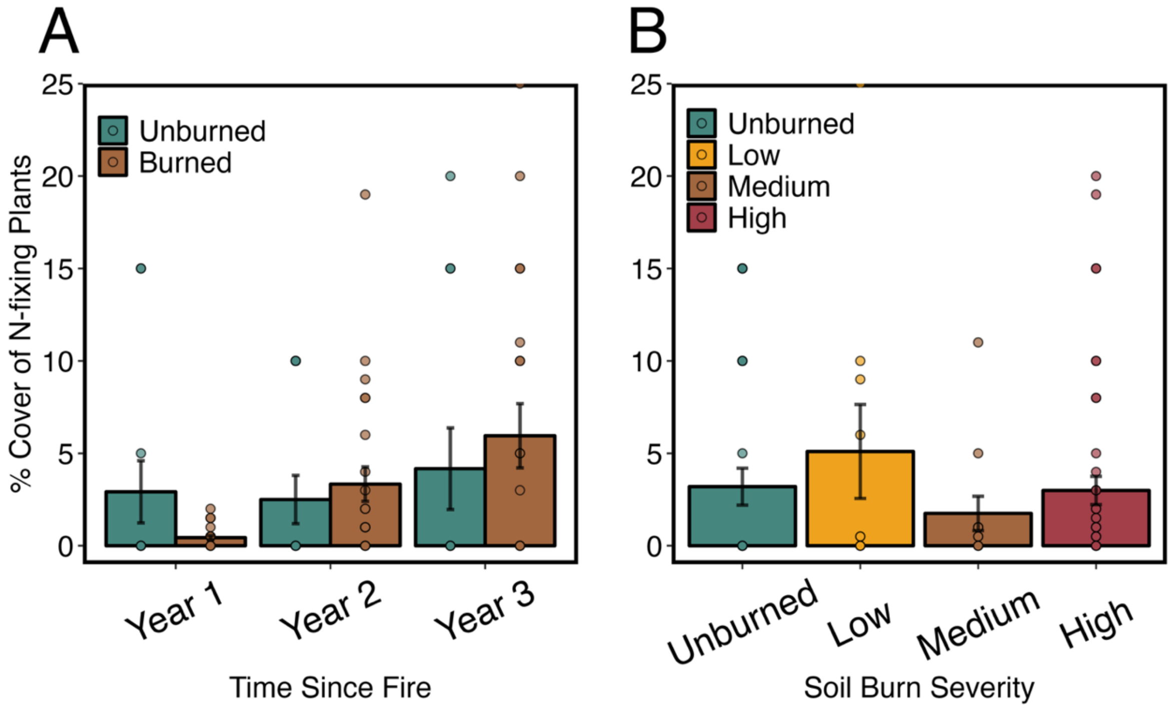


**Figure S6**. Mean % cover of N fixing species over each year of the study (**A**) and across each fine-scale (1 $m^2$) burn severity category (**B**). Dots represent individual measurements. Error bars are standard error, for each timepoint n = 12 unburned, n = 24 burned; plots that had no N-fixing species present were recorded as “0” and are therefore shown as overlapping data points at y = 0.

**Equations**

**Eq S1.**

```
NOmodel <- lme(
  logNO
~endWC+pH+C.N+MicroC+NH4_rate+Nitr_rate+Nfixer+PerClay+bac_mass+fun_mass,
  random = ~1 | PlotID,
  correlation = corAR1(form = ~ year | PlotID),
  weights = varIdent(form = ~1 | soil_type),
  data = data,
  method = "ML")
```

**Eq S2.**

```
N2Omodel <- lme(
  logN2O
~endWC+pH+C.N+MicroC+NH4_rate+Nitr_rate+Nfixer+PerClay+bac_mass+fun_mass,
  random = ~1 | PlotID,
  correlation = corAR1(form = ~ year | PlotID),
  weights = varIdent(form = ~1 | soil_type),
  data = data,
  method = "ML")
```

**Eq. S1, S2** – Equations used in our R script for global models of NO and $N_2O$**,** where logNO and log$N_2O$ = the log-transformation of cumulative NO and $N_2O$ over 40 h; endWC = gravimetric water content at the end of the gas flux incubation; C.N = bulk soil C:N ratio; MicroC = microbial biomass C (μg C g soil$^{-1}$); NH4_rate = net NH4 production rate during gas flux incubation; Nitr_rate = net nitrification rate during gas flux incubation; Nfixer = % cover of N-fixing plant species present at the plot; PerClay = % clay; and fun_mass and bac_mass = abundances of fungi (fun) and bacteria (bac).

### Supplementary Methods

#### *S1. Bacterial and Fungal Abundance and Richness*

We extracted DNA from 0.25g of sieved frozen (-80 °C) subsamples of the same soil cores used for soil chemistry monitoring and to measure NO and $N_2O$ fluxes with Qiagen DNeasy PowerSoil Kits. We measured bacterial and fungal gene copy number quantitative (q) PCR as a proxy for microbial abundance using the Eub338/Eub518 primers for bacteria (Fierer et al., 2005) and FungiQuant- F/FungiQuant- R primers for fungi (Liu et al., 2012). We also estimated microbial richness with Illumina MiSeq sequencing of the V4 region of the 16S rRNA gene with primer pair 515F and 806R (Caporaso et al., 2011) and fungal Internal Transcribed Spacer 2 (ITS2) with the primer pair ITS4-fun and 5.8 s (Taylor et al., 2016). Detailed methods for processing of qPCR and Illumina MiSeq data are provided in Pulido-Chavez et al. (2023).

#### *S2. Soil fluxes of NO and $N_2O$*

50 g of air-dried soils were wet to 75% WHC in 120-mL Mason jars and soil fluxes of NO and $N_2O$ were immediately measured for 48 h by connecting to a continuous-flow recirculating sample loop controlled by a multiplexer (LI-8150, LI-COR Biosciences) connected to a laser $N_2O$ analyzer (Los Gatos Research, Inc.; Model 914–0027), an infrared $CO_2$/$H_2O$ analyzer (LI-8100, LI-COR Biosciences), and a chemiluminescent $NO_2$ analyzer (Scintrex LMA-3D, Unisearch Associates, Canada) fitted with a $CrO_3$ converter to oxidize NO to $NO_2$ (when the $CrO_3$ converter was removed, $NO_2$ was not detectable, suggesting our measurements were mostly NO). Each jar was measured continuously for 13 minutes at intervals of 2.5 h. Because NO is consumed by the $NO_2$ analyzer, we used a 3-way solenoid valve to automatically route air from

the recirculating sample loop at a rate of 1.5 L $min^{-1}$ into the $NO_2$ analyzer halfway through the total 13-minute measurement cycle. At the same time, 1.5 L $min^{-1}$ zero-grade air (Ultra Grade Zero Air, Airgas, Radnor) was allowed to flow into the jars to replace the sample and prevent a vacuum. NO concentrations were allowed to equilibrate with the sample loop for 6.5 minutes and the last 30 s were averaged. Soil NO emissions were calculated as:

Eq. S3 $$\text{NO flux} = \frac{([\text{NO}]\text{outlet} - [\text{NO}]\text{inlet}) \times \text{flow } (1.5 \text{ L } min^{-1}) \times \text{g N}-\text{NO } (14 \text{ g } mol^{-1})}{\text{soil (g) x R x temp}}$$

where [NO]outlet is the concentration of NO leaving the jar headspace (ppb), [NO]inlet is the concentration of NO entering the jar (assumed to be 0 ppb), soil (g) is the mass of soil in the jar (50 g), R is the molar gas constant (0.0821 L atm $K^{-1}$ $mol^{-1}$), and temp is the room air temperature (~25 °C). At the end of the 13-minute measurement cycle, the sample loop was purged for 1 minute with ambient air before the next jar was measured.

$N_2O$ emissions were calculated as the increase in concentration over the first 6.5 minutes of the measurement cycle when the $NO_2$ analyzer was disconnected from the sample loop using a publicly available script (Andrews & Krichels, 2022). Trapezoidal integration (trapZ function in R; R Core Team, 2024) was used to calculate cumulative $N_2O$ and NO emissions over the 48-h incubation.

#### *S3. Natural abundance isotopic composition of $N_2O$*

To assess possible sources of $N_2O$ emissions, we measured isotopocules of $N_2O$ ($\delta^{15}N_2O_{bulk}$, $\delta N_2{}^{18}O_{bulk}$, and site preference (SP) $\delta^{15}N_2O_{SP}$, which reflects the placement of $^{15}N$ in the central (α) and peripheral (β) positions in the $N_2O$ molecule; where δ =

(((Rsample/Rstandard)−1)×1000) in units of ‰ and R = $^{15}N/^{14}N$ or $^{18}O/^{16}O$) in five burned soil samples that produced high $N_2O$ emissions and one unburned soil as reference. $N_2O$ was captured by incubating 50 g of soil wet up to 75% of water holding capacity in sealed air-tight 250-mL mason jars. Jars were fitted with 1-L gas-tight bags (Cali-5-Bond; Calibrated Instruments LCC) pre-filled with zero-grade air and the jar headspace was flushed with zero-grade air for 10 minutes. Jars were incubated at 25 °C for 24 hours, then the headspace was mixed by pumping with a 50 mL syringe 10 times (Stuchiner & von Fischer, 2022). The 1-L bags were sealed and analyzed within one week on a cavity ringdown infrared $N_2O$ analyzer (Los Gatos Research, Inc.; Model 914–0027) fitted with a Nafion water trap (PD-200 T-12MPS, Perma Pure LLC), a $CO_2$ trap (Carbosorb, Elemental Microanalysis), and an activated charcoal and silica gel (Sigma- Aldrich) trap to remove excess water and volatile organic compounds which can interfere with $N_2O$ isotopomer measurements (scrubber set-up described by Stuchiner et al., 2021; as published in Krichels et al., 2023). 1-L gas samples flowed into the isotopic $N_2O$ analyzer at a rate of 80 mL $min^{-1}$ to allow ~10 minutes of measurement time. Because it took 5–6 minutes for $N_2O$ concentrations to stabilize, $N_2O$ concentrations and $\delta^{15}N_2O_{SP}$, $\delta^{15}N_2O_{bulk}$, and $\delta N_2{}^{18}O_{bulk}$ were averaged over the last 3 minutes of our 1-s resolution data (n = 180) yielding ~± <0.002 standard deviations for all isotopocules concentrations in the range of our samples (**Table S6**; 1.6–2.8 ppm $N_2O$). Data were referenced to isotope standards United States Geological Survey (USGS)-51, USGS-52, USGS-32, USGS-34, and USGS-35 (Reston Stable Isotope Laboratory)  by applying individual standard curves ($R^2$>0.99; concentration range 1–7.5 ppm $N_2O$) for each isotopocule of $N_2O$ ($^{14}N^{15}N^{16}O$, $^{15}N^{14}N^{16}O$, and $^{14}N^{14}N^{18}O$). Isotopocule concentrations were converted to delta notation using equations derived from Stuchiner et al., (2021):

Eq.S4

$$\delta^{15}N^{\alpha} = \left[\frac{(N^{15}NO/N_2O)_{sample}}{(N^{15}NO/N_2O)_{standard}} - 1\right] * 1000$$

Eq. S5

$$\delta^{15}N^{\beta} = \left[\frac{\left({}^{15}NNO/N_2O\right)_{sample}}{\left({}^{15}NNO/N_2O\right)_{standard}} - 1\right] * 1000$$

Eq. S6

$$\delta^{18}O = \left[\frac{(NN^{18}O/N_2O)_{sample}}{(NN^{18}O/N_2O)_{standard}} - 1\right] * 1000$$

Site preference was calculated as:

Eq. S7

$$SP = \delta^{15}N^{\alpha} - \delta^{15}N^{\beta}$$

To reference our sample isotopic values to previously published ranges of isotopomers of $N_2O$ for distinct microbial processes such as nitrification (Ni), nitrifier denitrification (nD), bacterial denitrification (bD), and fungal denitrification (fD), we adjusted these values using the average combined soil $\delta^{15}N$-$NO_3^-$ and $\delta^{15}N$-$NO_2^-$ (substrates assumed to contribute to denitrification processes bD and fD) and $\delta^{18}O_{H2O}$ for nD, bD, and fD **(Table S5)** according to recommendations by Yu et al. (2020). Soil $\delta^{15}N$-$NO_3^-$ and $\delta^{15}N$-$NO_2^-$ were measured using the bacterial denitrifier method with *Pseudomonas aureofaciens* (Sigman et al., 2001). $\delta^{15}N$ and $\delta^{18}O$ values were measured using an isotope ratio mass spectrometer (Thermo Delta V, Thermo Fisher Scientific) and the Gas Bench interface. The international reference materials United States Geological Survey (USGS)-32, USGS-34, and USGS-35 were included in each analytical run and a value of -9 ‰ was used for $\delta^{18}O_{H2O}$ of deionized water in the building. We used the publicly available FRAME model to partition fractional contributions of $N_2O$ production processes (Ni, fD, nD, bD) and the residual $N_2O$ fraction (r) to estimate the proportion of $N_2O$ reduction (Lewicki et al., 2022; Well & Flessa, 2010; Yu et al., 2020).
